# Gut-derived metabolic reprogramming drives immune aging and tissue degeneration

**DOI:** 10.64898/2026.04.14.718497

**Authors:** Sayan Ghosh, Victoria Koontz, Ying Xin, Sridhar Bammidi, Dominique Meyer, Haochen Wang, Vishnu Suresh Babu, Puja Dutta, Durgadas Cherukaraveedu, Shreevadsaa A. Mohanakrishnan, Anupam K. Mondal, Jagannath Das, Jenny Nguyen, Avinash Soundararajan, Idris A. Adekale, Dulal Bhaumik, Stacey Hose, Sheldon Rowan, Padmanabhan P. Pattabiraman, Rangaramanujam M. Kannan, James T. Handa, Ji Yi, Srinivasa R. Sripathi, Jiang Qian, Debasish Sinha

## Abstract

Aging is characterized by changes in gut microbiome, metabolic imbalance and chronic inflammation, yet how these processes integrate to drive tissue degeneration remains poorly defined. Using age-related macular degeneration (AMD) as a model of tissue aging, we identify a diet-induced metabolic–immune axis that promotes systemic and retinal degeneration. In mice, a high-fat, cholesterol-enriched (HFC) diet induced perturbations in the gut structural integrity and microbiome repertoire, as well as systemic metabolic aging signatures, prominently marked by reduced circulating histidine. Plasma histidine levels were similarly decreased in AMD patients and inversely correlated with body mass index (BMI) in control donors. These diet-induced gut microbiome changes and subsequent metabolic alterations promoted peripheral innate immune reprogramming, with expansion of inflammatory neutrophils and monocytes that infiltrated the outer retina in a mouse model. Mechanistically, the gut-derived IGF1R/AKT2 signaling acts as a central regulator of global epigenetic remodeling and systemic immune aging under high-fat conditions in *C. elegans*. In a mouse model with an age-dependent dry AMD-like pathology, distinct retinal pigment epithelium (RPE) subpopulations exhibited downregulation of the histidine transporter SLC7A5, linking metabolic stress to activation of MIF/CD74-dependent inflammatory signaling between RPE and infiltrating immune cells. Histidine supplementation or AKT2 phospho-state modulation attenuated systemic immune activation and rescued retinal degeneration. These findings identify histidine-axis dysregulation as a mechanistic bridge between diet-induced microbiome changes, metabolic stress, immune aging, and retinal degeneration.

## Introduction

Aging involves the gradual accumulation of cellular and molecular damage that reduces tissue function and increases vulnerability to disease (1,2). Age-related macular degeneration (AMD), the leading cause of blindness among the elderly worldwide, exemplifies this process and serves as a powerful model of accelerated aging in post-mitotic tissue. It is now recognized as a systemic disease driven by metabolic dysfunction and chronic inflammation rather than a purely local retinal disorder (3,4). Previous studies have shown that the gut microbiome, metabolic dysfunction, and chronic inflammation are key contributors to this decline (5–7). The gut microbiome changes substantially with age (5), and these changes can influence peripheral metabolism such that alterations affect distant organs beyond the intestine (8–12), as evident in age-related diseases that show peripheral metabolic changes and chronic inflammation and immune dysregulation (13–27). However, the specific mechanisms whereby diet-induced gut microbial remodeling generates defined metabolic signals capable of reprogramming peripheral immune cells and converging on vulnerable tissues to drive degeneration remain poorly understood.

Nutrient sensing and amino acid metabolism are central to these aging processes (28,29). Several amino acids are known to decline with age in both humans and model organisms, and histidine is among the most consistently reduced (30–34). Low histidine levels correlate with increased inflammation, metabolic disease, and oxidative damage across multiple studies (35–39). Changes in amino acid availability can reprogram innate immune cells through a mechanism called trained immunity (40,41). These changes are often accompanied by epigenetic modifications that persist even after the initial trigger is removed (42). Trained immunity contributes to chronic inflammation during aging and contributes to several diseases (43–45). The gut microbiome directly influences circulating amino acid levels since microbial enzymes degrade dietary protein and synthesize amino acids that enter host circulation. In contrast, an increase in amino acid metabolizing bacteria in the gut may decrease the peripheral levels of those metabolites (46). Germ-free animals which lack a microbiome show altered amino acid profiles compared to conventionally raised animals (47,48). Antibiotic treatment that disrupts normal microbiome of the gut also changes plasma amino acid composition (49–51). Importantly, whether diet-driven alterations in gut microbiome and subsequent depletion of specific amino acids such as histidine activates defined molecular pathways to drive systemic innate immune reprogramming, and whether this process exacerbates degeneration in metabolically vulnerable tissues such as the retina, has not been eluded.

Across several model systems, metabolic and dietary interventions induce epigenetic changes that involve conserved longevity pathways including insulin and insulin-like growth factor 1 (IGF-1) signaling and AKT kinases, that are critically linked to maintenance of metabolic and inflammatory homeostasis (52–56). Specifically, AKT2 which is the downstream effector of IGF-1 has been implicated in epigenetic remodeling and inflammatory processes relevant to AMD pathogenesis (57). Interestingly, in the intestine, modulation of IGF1R/DAF-2 signaling extends lifespan and reduces systemic inflammation in *C. elegans* (53). However, whether gut-driven histidine depletion activates IGF1R/AKT2 signaling as a central node to drive systemic immune reprogramming through specific epigenetic mechanisms remains to be established.

Activated peripheral immune cells are pre-disposed to infiltrate tissues already under stress where they trigger chronic inflammation (58,59). The diseased retina shows accumulation of lipid-rich deposits, lysosome and mitochondria dysfunction, and display progressive cell loss along with many canonical hallmarks of chronic inflammation, such as immune cell infiltration and pro-inflammatory chemokines/cytokines activation (21–23,57,60). Substantial evidence in the literature links gut microbial composition to retinal aging and disease (13–15,61). Dietary interventions that reshape the microbiome can ameliorate or exacerbate retinal degeneration in animal models, and importantly, delay the progression of human diseases like age-related macular degeneration (AMD) (61–63), the most common cause of blindness among the elderly in the world (3). Despite these observations, the mechanisms by which gut alterations and amino acid perturbations affect retinal metabolism remain unknown. Additionally, whether systemic metabolic signals can reprogram the peripheral immune system and exacerbate retinal degenerative changes has not been verified.

Here, we delineate a pathway through which diet-induced gut microbial remodeling drives systemic aging by disrupting amino acid metabolism and reprogramming innate immunity. Using AMD as a model of accelerated aging in post-mitotic tissue, and employing a high fat, cholesterol-enriched (HFC) diet in C57BL/6J mice and the RPE-specific *Cryba1* conditional knockout (cKO) mouse model with age-dependent dry AMD-like pathology, alongside *C. elegans* as a genetically tractable system to interrogate conserved intestinal signaling, our data support a directional model in which HFC diet induced alterations in gut microbiome acts upstream of systemic histidine depletion, and systemic immune deregulation. We identified that gut derived IGF1R-AKT2 signaling pathway drives global epigenetic reprogramming, leading to predisposition to inflammatory activation. In parallel, age-dependent downregulation of SLC7A5 renders RPE cells metabolically vulnerable. The convergence of these systemic and tissue-intrinsic mechanisms culminates in MIF/CD74-dependent chronic inflammation and degeneration. These findings highlight therapeutic opportunities for age-related diseases driven by metabolic and immune dysfunction.

## Results

### Diet-induced gut microbiome remodeling is associated with systemic metabolic aging signatures

To investigate how dietary perturbations, affect the gut microbiome and systemic metabolism, we subjected 8-month-old WT C57bl6/j mice (male and female) to a high fat, cholesterol-enriched (HFC) diet regimen for 4 months (67). While high fat diet related gut microbiome changes are well documented, the impact of HFC, with distinct macro-nutrient composition and high cholesterol burden, remains unexplored. We employed a 16s rRNA sequencing-based assay on fecal samples from control and HFC fed mice. A total of 1,492,946 reads passed quality filters and chimera detection, leading to identification of 2,588 operational taxonomic units (OTUs). Adequate OTU identification in each sample was confirmed by rarefaction analysis. Principal component analysis (PCA) was performed on OTU abundances to observe global differences linked to diet treatment, which found distinct clustering of control and HFC microbiomes (Fig. 1a). Visual assessment suggested comparatively higher heterogeneity in the HFC gut microbiome profile when compared to the control. *Firmicutes* and *Bacteroidota* were most common, based on relative abundance at the phyla level, which showed typical profiles (Fig. 1b). An overlap of 461 taxa were present in both groups, while 443 and 41 were exclusive to HFC and control animals, respectively (Fig. 1c). These comparisons suggested an elevated taxonomic diversity in HFC animals that was confirmed through alpha diversity analysis (observed species) (Fig. 1d). Subsequently beta-diversity using weighted unifrac distance, which measures taxonomic diversity and abundance, revealed in-group clustering among control and HFC samples (Fig. 1e). To identify organisms significantly differentially represented in HFC gut microbiomes relative to control, linear discriminant analysis with LEfSe was performed and identified signature taxa at every taxonomic level (Fig. 1f). HFC microbiomes were enriched for *Clostridia*, *Proteobacteria* and *Desulfobacterota* among other taxa (Fig. 1g). At the same time, HFC samples had lower abundance of *Bacilli* and *Lactobacillales* (Fig. 1h). Overall, HFC induced large scale shifts in gut microbial compositions.

**Figure 1:**
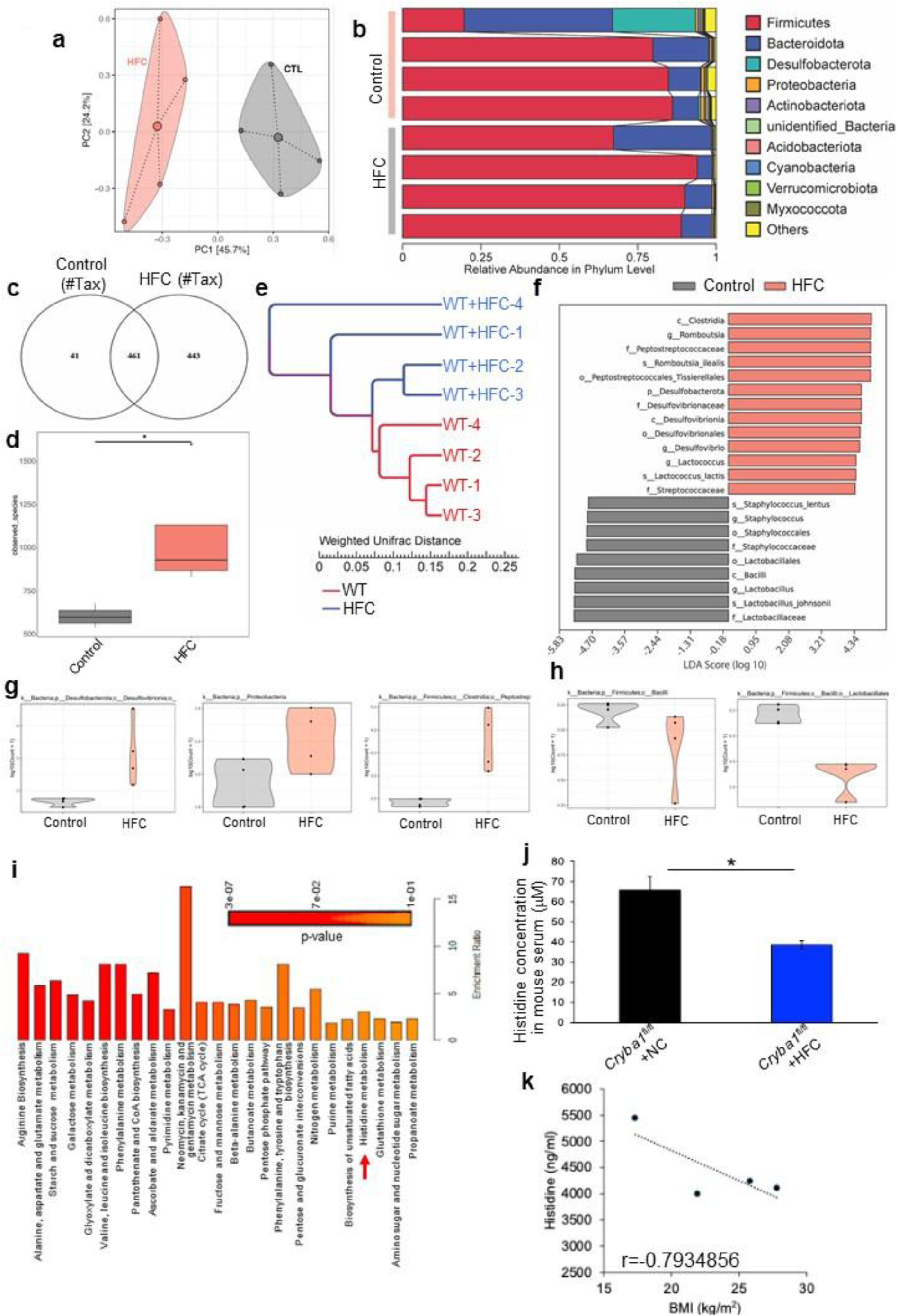
High fat diet modifies gut microbial communities in WT mice. (**a**) PCA biplot created with edgeR’s plotMDS shows gut microbiomes distributed over the first two principal components. (**b**) Relative abundance summaries at phylum level. (**c**) Venn diagram comparing taxa detection (group level) in the two diet groups. (**d**) Alpha diversity (observed_species) comparisons between control and HFC groups summarized in boxplots. (**e**) Dendrogram summary of weighted unifrac based relatedness among HFC and control microbiomes. (**f**) LEfSe plot showing LDA scores of significantly differential taxa between the diet groups. (**g**) Violin plots of HFC enriched taxa. (**h**) Violin plots of taxa depleted in HFC. Significantly enriched metabolic pathways represented as (**i**) over-representation analysis bar plot from serum metabolomics (GC-TOF MS) analysis in normal- and HFC-fed mice (arrow=significant enrichment of histidine metabolism pathway). n=4. (**j**) HPLC based quantitative estimation of histidine levels in mouse serum also showed similar downregulation in HFC-fed versus NC-fed floxed mice. n=3. (**k**) Correlation analysis showing inverse relationship between serum histidine levels and BMI among control donors. n=4. *P<0.05.

To determine whether gut microbiome alterations triggered systemic metabolic changes consistent with accelerated aging, we performed metabolomic profiling of peripheral blood. We found with pathway enrichment analysis by MetaboAnalyst 6.0 a significant enrichment in amino acid metabolism, particularly arginine biosynthesis, phenylalanine/tyrosine/tryptophan biosynthesis, alanine/aspartate/glutamate metabolism and histidine metabolism (arrow) pathways, known to decline with biological aging (Fig. 1i). Notably, circulating histidine levels were significantly reduced in HFC-exposed mice (Fig. 1j and Supplementary Fig. 1a). Histidine levels in the plasma/serum have been shown to decrease with age in multiple human cohorts independent of biological sex (28,29). To assess human disease relevance, we measured plasma histidine levels in dry/early AMD patients and age-matched healthy controls. AMD patients displayed lower histidine concentrations compared to controls with no AMD (Table-1 and Supplementary Fig. 1b). This finding is consistent with prior human metabolomics studies demonstrating decreased systemic histidine in late AMD (68) and perturbations in histidine-related metabolites in AMD plasma (69,70). Additionally, histidine levels inversely correlated with BMI in aged control donors (Fig.1k), implicating altered histidine metabolism in individuals who are metabolically stressed and with higher body weight (Table-1). Together, these studies support dysregulation of histidine metabolism as a reproducible metabolic feature of metabolic stress and in human AMD.

### Metabolic dysfunction drives peripheral immune aging through trained immunity

We next asked whether this metabolic shift associated with histidine depletion reprograms innate immunity. We hypothesized that amino acid depletion, particularly histidine deficiency, could influence peripheral innate immune reprogramming during AMD pathogenesis. Such reprogramming is typical of inflammaging, a chronic/sterile, low-grade inflammation that develops with advanced age and is independent of infections (71). To evaluate whether this metabolic perturbation following HFC feeding induced immune aging phenotypes associated with AMD, we performed comprehensive immune profiling in a mouse model with age-dependent retinal chronic inflammation, immune cell infiltration and a dry AMD-like phenotype (57,65,66). HFC feeding was started in *Cryba1* cKO mice at 8 months of age before onset of AMD-like phenotype, and continued for 4 months, well beyond onset of AMD-like changes (57,65,66). *Cryba1*^fl/fl^ (*Cryba1*-floxed) mice were used as controls. Neutrophil numbers were increased in peripheral blood of HFC mice using flow cytometric analysis (Supplementary Fig. 2 and Fig. 2a), a feature consistent with age-related chronic inflammation (72,73). In HFC-fed *Cryba1*^fl/fl^ (*Cryba1*-floxed controls) animals, these neutrophils also had elevated levels of CXCR2 (Fig. 2b), a resolution of inflammation marker (64). However, HFC-fed *Cryba1* cKO animals had lower CXCR2 levels when compared to normal chow fed *Cryba1* cKO or HFC-fed floxed controls, suggestive that neutrophil function was reprogrammed by HFC-induced metabolic stress (Fig. 2b), as has been reported previously in AMD patients (64). In addition, monocyte counts were also significantly elevated in HFC-exposed *Cryba1* cKO animals, as compared to normal chow fed *Cryba1* cKO or HFC-fed floxed (*Cryba1*^fl/fl^) controls (Fig. 2c). Interestingly, *Cryba1* cKO mice exhibited elevated basal expression of CCR2 in monocytes (Fig. 2d), indicative of systemic immune priming. CCR2 is a known pro-inflammatory marker associated with increased monocyte retinal infiltration in AMD (74). HFC independently increased the expression of CCR2 in floxed controls. However, no additive increase was observed in cKO mice under HFC conditions (Fig. 2d), recapitulating the development of a pro-inflammatory niche in the periphery and suggesting that age-driven inflammatory programming might predominate over dietary modulation. Furthermore, to assess if peripheral metabolic insults trigger immune activation in AMD, we assessed the inflammatory status of both neutrophils and monocytes *in vitro*. We found marked elevation of pro-inflammatory mediators (65,66), such as Lipocalin-2 (LCN-2), myeloperoxidase (MPO) and NETs (neutrophil extracellular trap) in cultured neutrophils from human donors (controls; with no eye disease) exposed to culture media supplemented with 10% human serum from AMD donors with the high risk *CFH* Y402H variant (H/H), compared to cells that were exposed to serum from controls without AMD and *CFH* Y402 (Y/Y) (Fig. 2e,f). Interestingly, histidine (10 mM) supplementation could reverse this activation (Fig. 2e,f). Additionally, we found elevated infiltration of CCR2+ cells in the outer retina of HFC-exposed cKO mice as well as in retinas from 12-month-old (aged) cKO mice, compared to floxed-controls fed with respective diets (Fig. 2g). These mice have been previously shown to have prominent infiltration of neutrophils Iba1-positive immune cells into the outer retina and sub-retinal space with age (65,66), suggestive of dysregulated immune homeostasis in the retina of this mouse model. Taken together, these findings suggest that gut alteration-driven histidine deficiency triggers systemic innate immune reprogramming and subsequent retinal infiltration, consistent with inflammaging, which could be associated with the exacerbation of several age-related retinal diseases like AMD.

**Figure 2:**
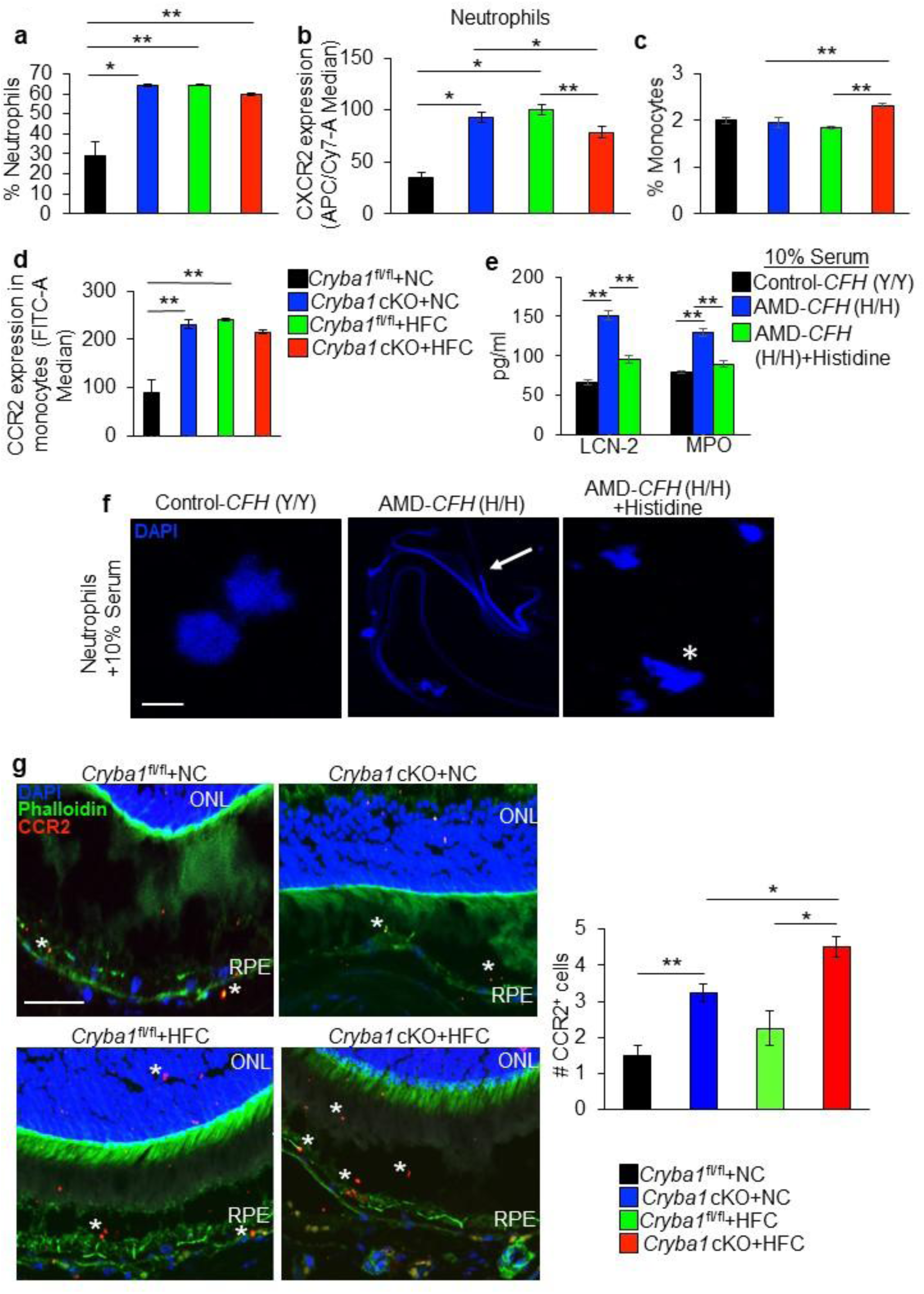
HFC-induced metabolic stress promotes peripheral immune activation and retinal infiltration. (**a**) Percentage increase of circulating neutrophils in normal chow (NC)-fed *Cryba1* cKO mice compared to floxed controls and HFC-fed animals from both genotypes compared to NC-fed mice. n=4. (**b**) CXCR2 (homeostatic marker) expression on circulating neutrophils showed decline in HFC-fed cKO mice, compared to HFC-fed *Cryba1*^fl/fl^ (floxed controls), whereas significantly increased expression was observed in HFC-fed floxed controls, relative to NC-fed mice of the same genotype n=4. (**c**) Circulating monocytes showed significant increase in HFC-fed cKO mice relative to HFC-fed floxed controls and NC-fed cKO animals. (**d**) The levels of CCR2 among monocytes also showed significant increase in HFC-fed animals from both genotypes and also in NC-fed cKO, compared to NC-fed floxed mice. n=4. (**e**) LCN-2 and MPO levels were increased in human neutrophils cultured with 10% serum from *CFH* Y402H AMD donors, compared to *CFH* Y402 (sibling controls; no AMD). Histidine (10 mM) supplementation reduced the levels of these proteins. n=4. (**f**) DAPI staining of neutrophils cultured with 10% serum from *CFH* Y402H AMD donors showed noticeable formation of NETs (arrow) compared to those cultured with serum from their *CFH* Y402 sibling controls (no AMD). This was rescued by histidine supplementation (asterisk). n=5. Scale bar=20 μm. (**g**) Representative immunofluorescence and quantification of outer retina stained for CCR2 (red; asterisks), phalloidin (green), and DAPI (blue), showed noticeable infiltration of CCR2+ cells in the outer retina, upon HFC exposure to floxed and cKO mice. n=4. Scale bar=50 μm. Data are mean ± SEM. *P<0.05, **P<0.01.

### High fat diet-induced intestinal dysfunction reprograms the gut-systemic immune axis

Given that histidine depletion likely originates from diet-induced microbiome alterations like increase in *Proteobacteria* and reduction in *Lactobacillus* (42), we next asked whether structural disruption of the intestinal barrier mediates this systemic immune reprogramming. We therefore examined intestinal barrier integrity, a critical determinant of systemic inflammation in aging (75). The distribution of E-cadherin, a key adherens junction protein responsible for intestinal barrier maintenance/integrity (75), was altered in the intestines of 12-month-old *Cryba1*-floxed mice under HFC conditions compared to normal chow as visualized with immunofluorescence confocal microscopy (Fig. 3a). We also observed severe disruption of villus architecture and reduced intestinal villi length in HFC fed *Cryba1*-floxed and cKO mice compared to mice fed normal chow, indicating compromised barrier integrity (Fig. 3b). To ascertain that gut structural alterations drives peripheral immune changes, we fed HFC to 5-month-old wild-type (WT) mice for 2 months with or without fecal microbiota transplantation (FMT) from normal chow fed littermates (Fig. 3c). Fecal transplantation rescued gut structural changes after HFC exposure, as evident from maintenance of villi structure (arrow in Fig. 3d). Furthermore, flow cytometry analysis showed that FMT reduced the levels of CD45^high^CD11b^high^Ly6C+Ly6G^high^ neutrophils in peripheral blood when compared to samples from mice fed HFC alone (Supplementary Fig. 3, Fig. 3e). Interestingly, FMT had no impact on the levels of inflammatory monocytes and monocytic myeloid-derived suppressor cells (M-MDSCs) (CD45^high^CD11b^high^Ly6C^+^Ly6G^low^) (Fig. 3f). This likely indicates an enhanced, yet potentially protective, immune response where the transplant actively modulates inflammation induced by the HFC rather than simply suppressing it. Thus, it is possible that rescuing gut homeostasis under metabolic stress rescues peripheral immune alterations (76). These findings suggest that maintaining gut homeostasis is critical for systemic immune regulation.

**Figure 3:**
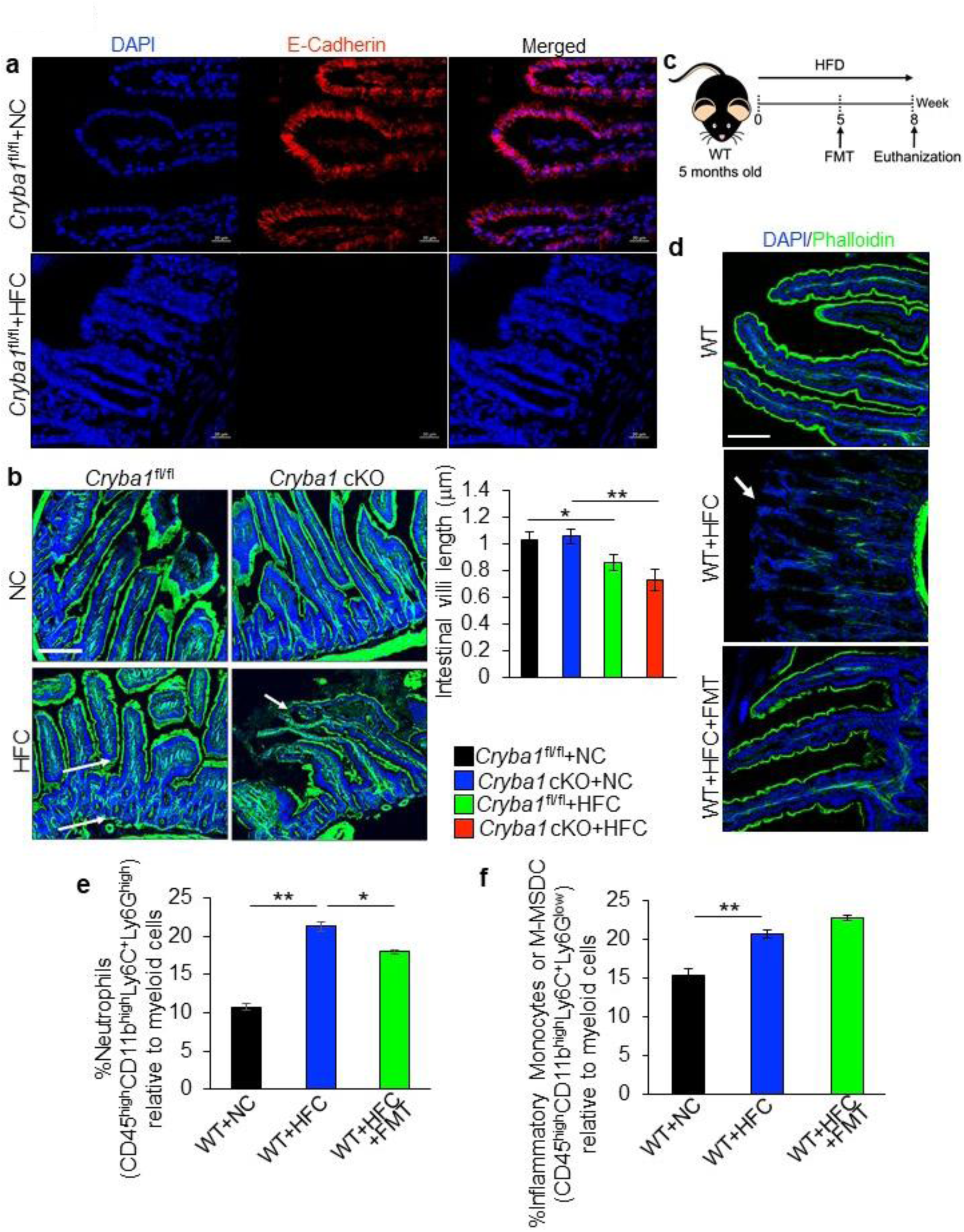
Intestinal dysfunction links HFC exposure to systemic immune remodeling. **(a)** Representative intestinal sections stained for E-cadherin and DAPI from *Cryba1*^fl/fl^ mice fed NC or HFC, showing robust downregulation of E-Cadherin (red) in HFC-diet. n=4. Scale bar=20 μm. (**b**) Representative immunofluorescence images (Phalloidin: Green) of intestinal sections with quantification of villus length from *Cryba1*-floxed and cKO mice fed with NC or HFC, showing marked alterations (arrow) and decrease in villi length upon HFC exposure. n=4. Scale bar=20 μm. (**c**) Schematic showing fecal microbiota transplantation (FMT) experimental design. (**d**) Representative intestinal morphology in WT, WT+HFC, and WT+HFC+FMT groups stained with E-Cadherin (red) and phalloidin (green) showing protection of intestinal structure upon FMT treatment compared to HFC-exposed mice without FMT (arrow) n=4. Scale bar=20 μm. (**e**) Percentage of circulating neutrophils (CD45^high^CD11b^high^Ly6C^+^Ly6G^high^) and (**f**) inflammatory monocytes/M-MDSCs (CD45^high^CD11b^high^Ly6C^+^Ly6G^low^) relative to myeloid cells showed significant and expected upregulation in HFC-exposed mice, compared to normal chow fed animals. FMT treatment reduced the levels of circulating (**e**) neutrophils and showed (**f**) relative increase in the levels of inflammatory monocytes/M-MDSCs, compared to HFC-exposed mice without FMT. n=3. Data are mean ± SEM. *P<0.05, **P<0.01.

### Gut-associated IGF1R signaling mediates epigenetic changes associated with accelerated immune aging

The involvement of gut health has been critically associated with aging and age-related detrimental effects both in humans and other species that share numerous conserved genetic and molecular pathways (53,54). To further validate the role of intestinal homeostasis in systemic inflammation, we focused on the evolutionarily conserved insulin/IGF-1 signaling (IIS) pathway using an established model to study gut function. In the intestine-specific inducible Daf2/IGF1R *C. elegans* line (daf-2(hq363[daf-2::degron::mNeonGreen]) III; *Daf2*^ins^), intestinal-specific knockout *Daf2* increases longevity and confers healthy aging in distant tissues (53). Synchronized *Daf2*^ins^ worms that were given high fat (HF) food for 72 h and treated with auxin (1 mM) to specifically knockout *Daf2* in the intestine, which resulted in a noticeable decrease in gene expression of the critical pro-inflammatory mediator, *Mek2* (mammalian MEK1/2) and *Tir-1* (mammalian SARM1; Sterile Alpha and TIR Motif-containing 1), compared to HF-fed *Daf2*^ins^ worms not treated with auxin (Fig. 4a,b), suggesting gut-derived *Daf2* signaling triggers acute systemic inflammation during metabolic stress. Histidine supplementation (10 mM) rescued the levels of these genes in HF-exposed *Daf2*^ins^ worms (Fig. 4a,b).

**Figure 4:**
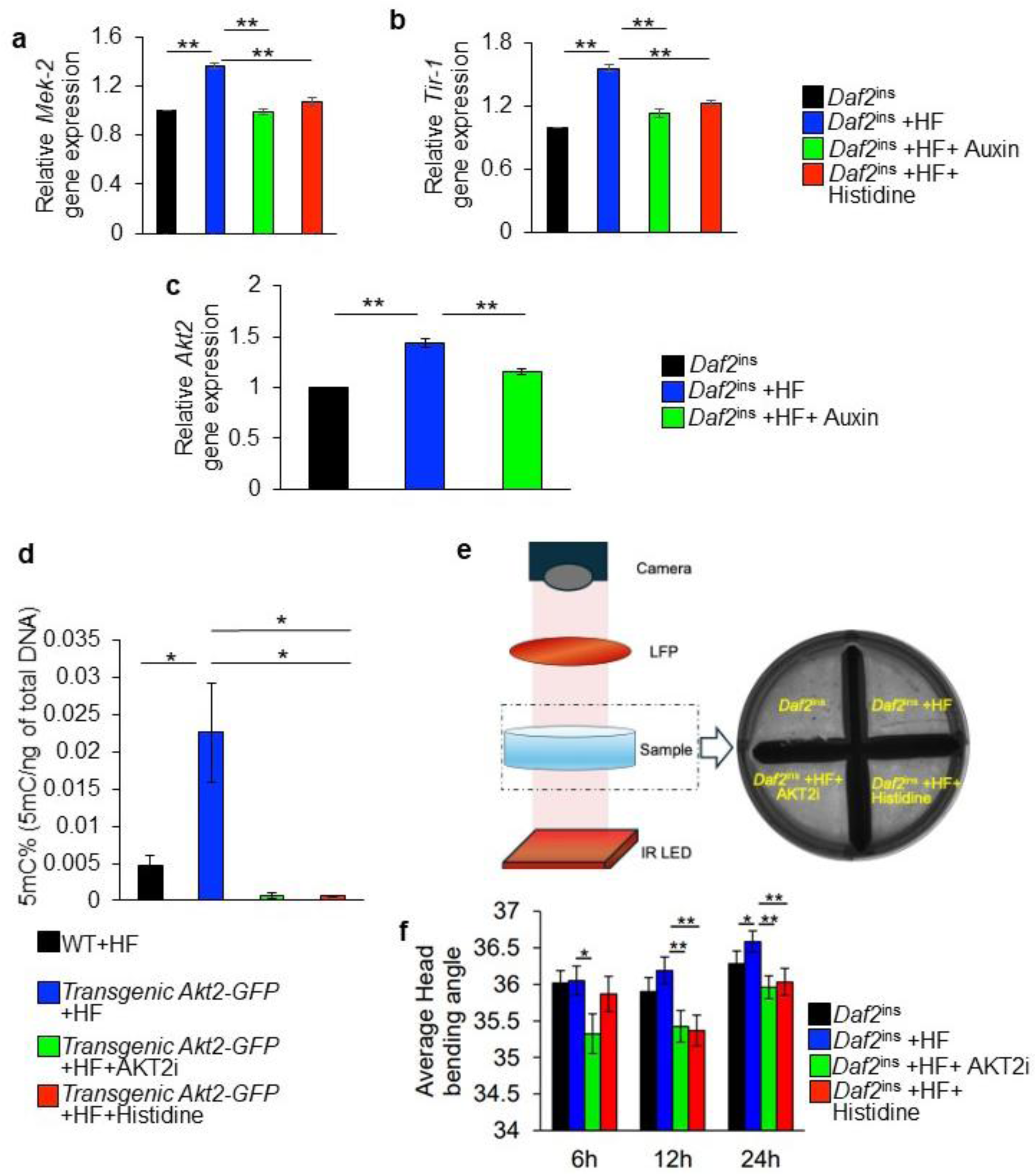
IGF1R/AKT2 signaling regulates inflammatory and epigenetic remodeling under metabolic stress. PCR analysis in synchronized *C. elegans* showed significant increase in the gene expression of immune regulators like (**a**) *Mek-2* and (**b**) *Tir-1* upon high fat (HF) exposure for 72 h, which was reduced when auxin (1 mM) was added to degrade intestinal *Daf2* specifically or by histidine supplementation (10 mM). n=3. (**c**) *Akt2* gene expression in *Daf2*^ins^ *C. elegans* following HF exposure was markedly increased, but was reduced significantly upon treatment with auxin (1 mM). n=3. (**d**) Global DNA methylation (5mC) levels were significantly elevated in transgenic *Akt2*-GFP worms exposed to HF, compared to WT+HF worms, which were rescued upon AKT2-PSM (AKT2i) or histidine treatment. (WT+HF and transgenic *Akt2*-GFP+HF; n=7, transgenic *Akt2*-GFP+HF+AKT2i and transgenic *Akt2*-GFP+HF+histidine; n=4). (**e**) Schematic of the health span assay, which was performed in a single well of a six-well plate. The plate is illuminated from below with an infrared (IR) LED array. Transmitted light passes through a long-pass filter (LPF) before being captured by a camera. The zoomed-in view shows the camera field of view, in which each well is subdivided into four regions, allowing four experimental conditions to be recorded simultaneously. After 24h, (**f**) head angle bending is noticeably increasing in HF-exposed *Daf2*^ins^ worms which was reduced upon AKT2-PSM (AKT2i) and histidine supplementation. n=3. Data are mean ± SEM. *P<0.05, **P<0.01.

One downstream effector of IGF1R/*Daf2* is AKT2, a key mediator of epigenetic and inflammatory processes (57,65,66,77). We found that high fat exposure triggered *akt-2* gene upregulation in *Daf2*^ins^ worms which was rescued by auxin treatment (Fig. 4c). To further corroborate the influence of AKT2 signaling on epigenetic changes, we performed global epigenetic profiling in worms with an additional extrachromosomal copy of AKT2 (mgEx341 [akt-2::GFP::unc-54 3’UTR + rol-6(su1006)]; transgenic *Akt2-GFP*). Our data showed global increase in DNA methylation (5-methylcytosine, 5mC) in transgenic *Akt2-GFP* worms exposed to HF (Fig. 4d), compared to WT worms exposed to HF. To establish causality of AKT2 signaling with histidine metabolism, we treated these worms with an AKT2 phospho-state modulator (AKT2-PSM; AKT2i at 10 nM) (78) or with histidine supplementation (10 mM) and observed that both treatments prevented HF-induced epigenetic remodeling, with DNA methylation significantly downregulated (Fig. 4d). Additionally, head bending angle was measured as a functional readout of physiological state, since changes in posture reflect underlying metabolic activity, fat accumulation, and neuromuscular health in worms. In *Daf2*^ins^ worms exposed to HF, the head bending angle increased at 24 hours, indicating a wider and more rigid posture consistent with elevated adiposity and metabolic impairment. This effect was significantly reduced by AKT2-PSM (AKT2i) or histidine supplementation (Fig. 4e,f and Movie-1). These findings suggest that HF stress induces a low-metabolic, developmentally arrested state resembling dauer, while AKT2 inhibition or histidine helps restore an active physiological condition. These findings suggest that gut derived IGF1R/AKT2 signaling is a master regulator of global epigenetic changes and that if perturbed, can accelerate metabolic-immune aging.

### Tissue-specific manifestation of metabolic changes and chronic inflammation in the retina

We next asked whether systemic immune reprogramming converges with intrinsic metabolic retinal vulnerability. The retina, with its unusually high metabolic demand and limited regenerative capacity, serves as a sensitive readout of organism-wide aging (79). Single-cell RNA sequencing (scRNAseq) of RPE/choroid tissue from aged (15-month-old) WT and *Cryba1* cKO mice (65), showed heterogenous transcriptomic RPE cell sub-populations characterized by differences in inflammatory and metabolic genes/pathway signatures between these populations (Fig. 5a,b). Our data also shows multiple infiltrating immune cells like neutrophils, monocytes, and macrophages, which has been shown to be critical in AMD pathogenesis (21,65,66,74), and each of which exhibited activation signatures characteristic of immune dysregulation, including noticeable enrichment of several metabolic and immune-modulatory/pro-inflammatory signaling pathways Fig. 5a and Supplementary Fig.4a-f). Additionally, Qiagen Ingenuity Pathway Analysis (IPA) comparative bioinformatic analysis between mouse RPE clusters and iPSC-RPE cells from *CFH* Y402H AMD donors were performed. Our analysis showed enrichment of similar inflammatory (green arrows) and metabolic (red arrows) genes/pathways (Fig. 5c), suggestive that transcriptional signatures accelerate cellular aging in the diseased state both in the mouse model and humans. Interestingly, an altered Class 1 MHC-dependent antigen presentation gene cluster was enriched in both cKO and AMD RPE (Fig. 5b,c), as were pathways for eukaryotic protein synthesis pathways, oxidative phosphorylation and other metabolic pathways, consistent with elevated mTORC1 signaling that has been shown previously in both the mouse model and human AMD RPE (80,81). Furthermore, following comprehensive differential gene expression analysis across multiple cell types from RPE-choroid of 3- and 15-months old WT and *Cryba1* cKO, *Slc7a5* expression was notably altered in specific RPE subpopulations (specifically clusters 16 and 27) and in immune cells from *Cryba1* cKO compared to controls at both 3 and 15 months (Fig. 5d). We also found that in human datasets of scRNAseq of the retina that were analyzed by the spectacle database software, that *Slc7a5* gene expression is noticeably downregulated in the RPE with increasing age and in both dry and neovascular AMD, compared to controls (Supplementary Fig. 5a,b). Furthermore, SLC7A5 immunolabeling was decreased in the RPE of *Cryba1* cKO and HFC treated cKO and floxed mice as well as human AMD retina (Supplementary Fig. 6a,b). Mechanistically, ChIPseq analysis (82) further indicated the causal link between histidine deregulation and RPE alterations. H3K27ac ChIP-seq profiles at the *Slc7a5* locus, a gene responsible for histidine transport and mTORC1 regulation, was striking reduced in histone acetylation signal within an intronic region (chr8:121,895,526–121,895,926) in RPE cells from 4-month-old *Cryba1* conditional knockout (cKO) mice, compared to floxed controls. This region shows clear H3K27ac enrichment in both FL replicates, indicating active enhancer or regulatory function under normal conditions. In contrast, this peak is notably diminished or absent in cKO RPE, as shown in both individual and combined (*Cryba1*^fl/fl^ and cKO) track views (Fig. 5e), suggesting decreased gene expression that mirrors predisposition to altered histidine metabolism in cKO RPE cells. These results support the model that aging triggers downregulation of *Slc7a5* in the RPE with the resultant metabolic stress that triggers detrimental signaling pathways and disease onset.

**Figure 5:**
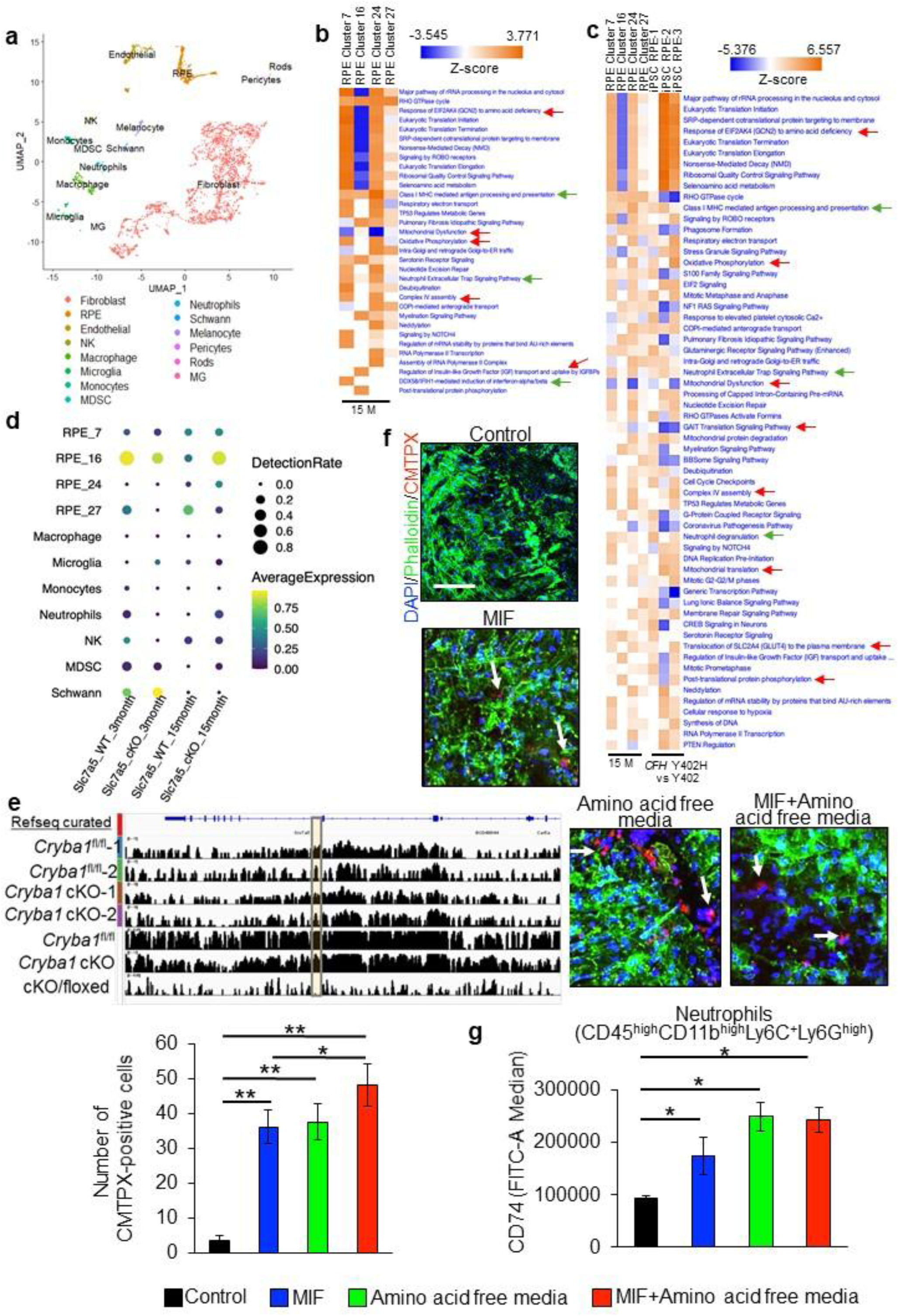
*Cryba1* deficiency triggers predisposition to metabolic and inflammatory insults in the RPE. (**a**) UMAP (Uniform Manifold Approximation and Projection) plot from scRNAseq of RPE/choroid isolated from aged (15-month-old) WT and *Cryba1* cKO mice reveals heterogeneous RPE subpopulations as well as several immune and other cell types. n=3. Heat map showing pathway enrichment in each (**b**) RPE sub-populations and (**c**) cross-species comparative transcriptomic analysis between mouse *Cryba1* cKO RPE clusters and human iPSC-derived RPE from *CFH* Y402H AMD donors. The enrichment showing inflammatory (green arrows) and metabolic (red arrows) pathways, indicating transcriptional programs suggestive of accelerated cellular aging in both *Cryba1*-deficient mouse RPE and human AMD RPE. n=3. (**d**) Integrated scRNAseq analysis of RPE/choroid from 3- and 15-month-old WT and *Cryba1* cKO mice demonstrating differential *Slc7a5* expression across cell populations. *Slc7a5* gene expression is significantly altered in specific RPE clusters (clusters 16 and 27) in *Cryba1*-floxed and cKO mice at both ages, indicating persistent histidine metabolism reprogramming during aging and disease progression. n=3. (**e**) H3K27ac ChIP-seq tracks at the *Slc7a5* (yellow highlighted inset) locus (chr8:121,895,526–121,895,926) in RPE from 4-months-old *Cryba1*^fl/fl^ and cKO mice. Floxed replicates show prominent H3K27ac enrichment within an intronic regulatory region, indicative of active enhancer activity. This acetylation peak is markedly reduced in cKO RPE (individual and combined track views), consistent with decreased chromatin accessibility and reduced Slc7a5 expression, linking *Cryba1* loss to histidine transport and mTORC1 dysregulation. (**f**) Functional RPE-mediated immune clearance assay was performed, after pre-treatment for 6 hours with recombinant MIF (1 ng), amino acid free media, or MIF+amino acid free media, followed by staining these cells with CMTPX-red and then co-culturing them with WT RPE explants for 24 hours. Quantification of residual CMTPX-positive cells demonstrates significantly reduced immune cell clearance following MIF and amino acid free media treatment, which was maximum when immune cells were exposed to MIF+amino acid free media, compared to untreated controls, indicating impaired RPE-mediated immune resolution under nutrient stress. n=4. (**g**) Flow cytometric analysis of treated immune cells showing selective upregulation of CD74 (FITC-A) in CD45^high^CD11b^high^Ly6C^+^Ly6G^high^ neutrophils following MIF, amino acid deprivation, or combined treatment. n=3. Data are mean ± SEM. *P<0.05, **P<0.01.

Our single cell sequencing data implicated amino acid transport and metabolism in RPE dysfunction during AMD pathogenesis. To ascertain the effect of *Cryba1* loss, which impairs lysosome function, on RPE metabolic changes, we performed metabolomic profiling in the RPE. Our results revealed extensive enrichment in several amino acid metabolic pathways in *Cryba1* cKO and floxed-control RPE (Supplementary Fig. 7a). Lipid biosynthetic pathways are strongly activated by mTORC1 signaling, which is regulated at the lysosome. We found major lipid metabolism genes in WT RPE overexpressing βA3/A1-crystallin or either of its two crystallin isoforms (βA1 and βA3), significantly downregulated compared to control WT RPE explants (Supplementary Fig. 7b). These data further support the importance of *Cryba1* in RPE metabolic homeostasis during aging and AMD pathogenesis.

Chronic retinal inflammation in AMD is governed by activation of proinflammatory mediators and subsequent infiltration of immune cells (21,65,66,74). We have previously shown significant sub-retinal infiltration of microglia and neutrophils in *Cryba1* cKO mice with age (65,66). We therefore investigated if RPE-specific molecular predispositions and simultaneous peripheral immune alterations triggered by HFC exposure, could drive chronic inflammation and exacerbate AMD-like pathology in the retina. Our scRNA-seq pathway analysis revealed noticeable enrichment of several inflammatory pathways in both cKO and human AMD RPE. Using comprehensive ligand-receptor (LR) interaction analysis using single-cell transcriptomic data of RPE/choroid tissue of WT and *Cryba1* cKO mice (65), we explored the cellular interactions driving chronic retinal inflammation at 3- and 15-months age (Supplementary Fig. 8a,b). Computational analysis revealed two different RPE sub-populations/clusters (RPE-16 and RPE-27) that had downregulated expression of *Slc7a5* and elevated expression of inflammatory mediators including MIF (Macrophage Migration Inhibitory Factor), APP (Amyloid Precursor Protein), APOE (Apolipoprotein E), A2M (Alpha-2-Macroglobulin), and THBS1 (Thrombospondin-1). Conversely, infiltrating immune cells like neutrophils, MDSCs and macrophages showed significant expression of their cognate receptors (Supplementary Fig. 8a,b). Importantly, ligand-receptor pairing identified a MIF/CD74 and APOE/CD74 signaling axis as a potential bridge between RPE metabolic stress and immune activation. Aged (where AMD-like phenotype is evident), stressed RPE cells, particularly RPE-16 and 27 clusters with lower levels of *Slc7a5*, expressed elevated levels of MIF and APOE while resident immune cell, microglia and infiltrating immune cells like neutrophils, macrophages, and MDSCs expressed the CD74 receptor (Supplementary Fig. 8a,b). This temporal analysis identified progressive expansion of this inflammatory network with age. Specifically, at 15 months, the cellular communication landscape intensified with recruitment of MDSCs expressing complement component C3, fibronectin (FN1), and immunosuppressive TGFB1 (Supplementary Fig. 8a,b). Notably, immune cells expressed matrix remodeling factors including MMP9 and SERPINE1, while widespread TGFB1 signaling suggested attempted, but failed, inflammation resolution (Supplementary Fig. 8a,b). This predicted MIF/CD74 axis likely represents a critical signaling hub for chronic inflammatory transitions in AMD pathogenesis and a druggable therapeutic target for interrupting the RPE-immune inflammatory circuit.

RPE cells are normally anti-inflammatory and effective at nullifying infiltrating immune cells in the sub-retinal space (SRS), and its failure has been shown to be critical for eliciting chronic inflammation during AMD pathogenesis (21–25,65,66,74,83). Therefore, to understand the role of the MIF/CD74 signaling axis in inflammation and exacerbation of RPE damage during nutrient stress and AMD, we exposed mouse peripheral immune cells with either: (i) MIF protein; (ii) amino acid-free media (to mimic systemic depletion of amino acids like histidine); (iii) MIF+amino acid free-media (to mimic both systemic and RPE-mediated activation cues) for 6 hours and then stained these cells with cell tracker dye, CMTPX followed by co-culture with 4-month-old mouse WT RPE explants for 24h. The percentage of immune cell clearance by the RPE was measured by counting CMTPX-positive cells on WT RPE explant cultures. We found that immune cells exposed to MIF, amino acid-free media or MIF+amino acid-free media were cleared less efficiently by RPE cells as compared to untreated immune cells (Fig. 5f). Flow cytometry analysis of these cultured immune cells also revealed that MIF only treatment, amino acid free media, and MIF+amino acid free media treatment all lead to significant upregulation of CD74 expression in CD45^high^CD11b^high^Ly6C^+^Ly6G^high^ neutrophils (Fig. 5g and Supplementary Fig. 9), but not in CD45^high^CD11b^high^Ly6C^+^Ly6G^low^ inflammatory monocytes or M-MDSCs (data not shown), thereby suggesting that neutrophil but not monocytes or M-MDSCs might be responsible in immune dysregulation and inflammation during metabolic stress. This further corroborated the scRNAseq LR loop analysis that showed enriched MIF/CD74 signaling in neutrophils and macrophages, but not in monocytes from aged *Cryba1* cKO RPE/choroid tissue (Supplementary Fig. 8).

Together, these findings suggest that SLC7A5-mediated amino acid transport dysfunction triggers a biphasic cascade in which early RPE predisposition leads to age- or stress-induced metabolic crisis, failed compensation, and ultimately chronic MIF/CD74-driven inflammation. These findings also suggest that systemic metabolic–immune aging processes converge in the retina, promoting sustained inflammation as a feature of accelerated tissue aging.

### Metabolic intervention rescues retinal degeneration in an acute model of dry AMD

Having shown that gut microbiome changes was associated with histidine deficiency which in turn drives the metabolic-immune aging cascade, and that supplementation with histidine can rescue peripheral immune changes, we wanted to ascertain if the metabolic abnormalities and immune cell infiltration correlated with noticeable exacerbation of RPE damage. We showed an increased number of dysmorphic RPE cells following HFC exposure in the 12-months-old *Cryba1* cKO mice that were fed with HFC for 4 months (Fig. 6a). We assessed the potential for metabolic intervention in an independent and non-genetic model of oxidative stress, a known contributor to AMD, caused by injection of sodium iodate (84), which stresses the RPE and causes retinal degeneration (Supplementary Fig. 10) in mice fed a HFC diet (Fig. 6b). Histidine supplementation or AKT2-PSM (AKT2i) treatment noticeably protected photoreceptor (rhodopsin loss; arrow) and RPE degenerative changes (asterisk), as evident from decline in the levels of Ezrin-Radixin-Moesin-binding phosphoprotein 50 (EBP50, marker of RPE polarity which is reduced during early/dry AMD pathogenesis) (57) in HFC+NaIO3 treated mice (Fig. 6c). Interestingly, AKT2i treatment after HFC+NaIO_3_ exposure effectively reduced HFC-induced elevation of peripheral immune cell populations, previously implicated in AMD (61,65,66,74), including levels of CD45^high^CD11b^high^Ly6C^+^Ly6G^high^ neutrophils and CX3CR1^+^ mononuclear phagocytes (CD45^high^Ly6G^-^F4/80^+^) (Fig. 6d,e and Supplementary Fig. 3,11). Additionally, histidine supplementation similarly rescued CX3CR1^+^ mononuclear phagocyte numbers, but showed only modest decrease, rather than significant rescue in neutrophil levels (Fig. 6d,e and Supplementary Fig. 3,11). NaIO_3_ alone diminished the levels of these immune cells (Fig. 6d,e and Supplementary Fig. 3,11) since elevated concentrations of sodium salts in the bloodstream can directly cause cytotoxicity to peripheral white blood cells (85). Therefore, the significantly increased levels of neutrophils and mononuclear phagocytes in the HFC+NaIO_3_ group relative to WT and NaIO_3_ only treatment, is likely solely due to the HFC treatment (Fig. 6d,e and Supplementary Fig. 3,11). These results indicate that targeting metabolic/inflammatory pathways involving histidine and AKT2 might be crucial in reducing peripheral immune activation even under acute stress. Collectively, these findings indicate that histidine supplementation is a potent metabolic intervention that reverses systemic aging phenotypes and prevents immune alterations, thus validating histidine metabolism as a central, modifiable regulator of biological aging and age-related diseases like AMD (Fig. 6f).

**Figure 6:**
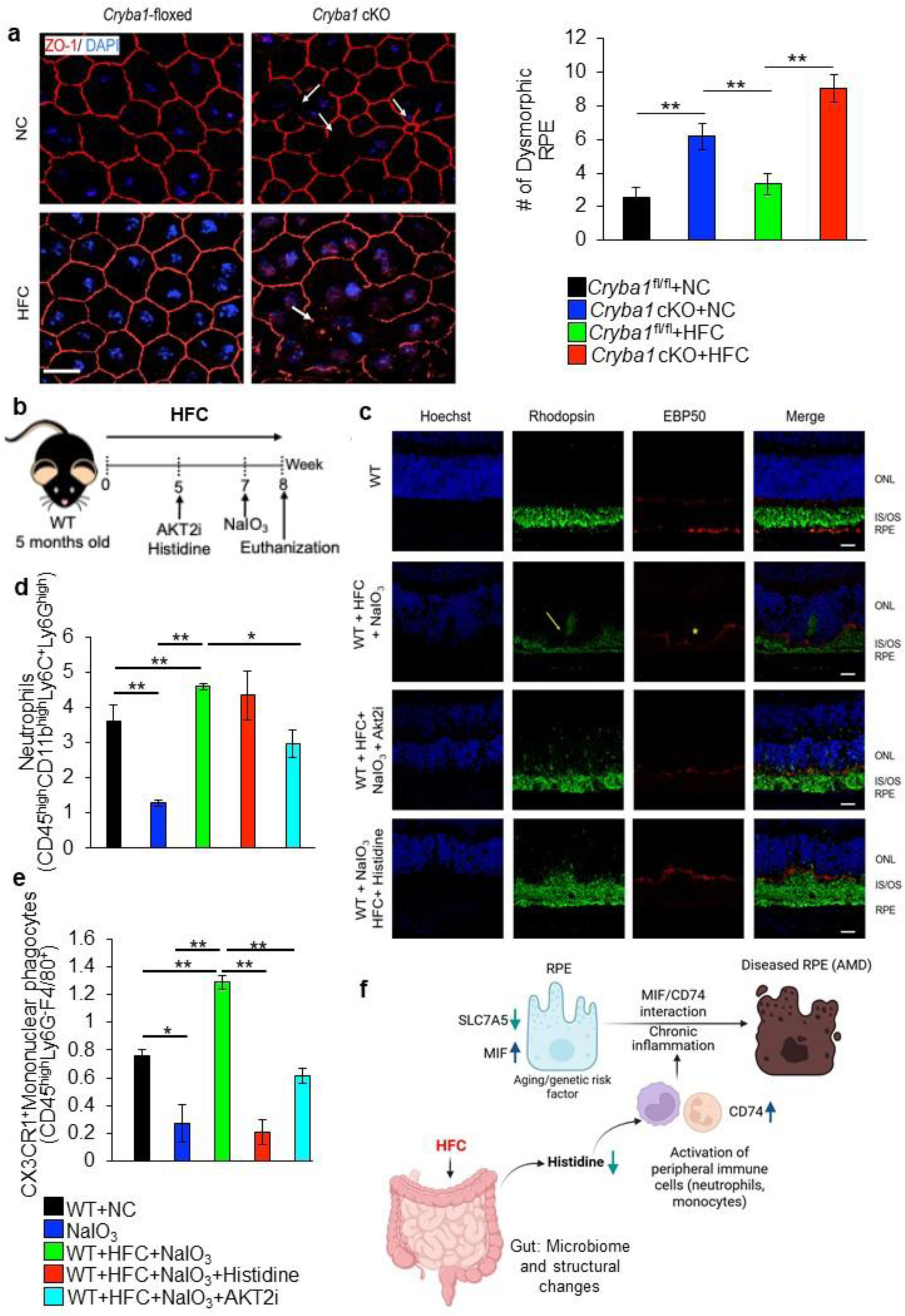
Targeting systemic AKT2 and histidine signaling could potentially reduce HFC-mediated peripheral immune activation and exacerbation of retinal degeneration. (**a**) Quantification of dysmorphic RPE (arrows) in *Cryba1*^fl/fl^ and cKO RPE flat mounts from mice fed with either NC or HFC. HFC exposure significantly increases number of dysmorphic RPE mice in cKO mice, indicating exacerbation of structural degeneration under metabolic stress. Such changes were not visualized in HFC-fed floxed mice. n=4. (**b**) Experimental schematic of acute dry AMD model induced by NaIO₃ under HFC conditions with histidine supplementation or AKT2-PSM (AKT2i) therapy. (**c**) Representative immunohistochemical images of outer retinal layers following NaIO₃ treatment with or without HFC exposure. NaIO3 and HFC+NaIO₃ induces photoreceptor degeneration (rhodopsin loss; arrows) and RPE damage as evident from reduced EBP50 expression (asterisk). Histidine supplementation or AKT2-PSM (AKT2i) therapy in HFC+NaIO₃ treated mice rescued these anomalies. n=3. Scale bar= 20 μm. Flow cytometric quantification of (**d**) CD45^high^CD11b^high^Ly6C^+^Ly6G^high^ neutrophils and (**e**) CX3CR1^+^ mononuclear phagocytes (CD45^high^Ly6G^-^F4/80^+^) in peripheral blood showing that HFC+NaIO₃ significantly increases neutrophil levels compared to WT and NaIO₃-only groups, indicating diet-driven immune activation. AKT2-PSM (AKT2i) therapy effectively restores neutrophil levels toward baseline, whereas histidine supplementation produces a relative but not statistically significant reduction. n=3. (**f**) Schematic showing that HFC treatment triggered gut microbiome and structural alterations, peripheral metabolic (decrease in histidine levels) and immune alterations with upregulated CD74 expression in neutrophils. These cells are able to infiltrate the retina and interact with RPE-derived MIF triggering a chronic inflammation and subsequent retinal degeneration as seen in AMD. Data are mean ± SEM. *P<0.05, **P<0.01.

## Discussion

AMD is increasingly recognized as a systemic disease influenced by metabolic dysfunction and chronic inflammation rather than a purely local retinal disorder (13–15,21–23). Aging is characterized by gut microbiome alterations, inflammaging, and metabolic decline (1,2,5), yet direct mechanistic links between diet-induced metabolic stress and retinal degeneration have remained unclear. Our study identifies dysregulation of histidine metabolism as a central metabolic mediator that connects HFC-induced gut dysfunction to systemic immune reprogramming and retinal pathology. By integrating microbiome, metabolomic, immunologic, and single-cell transcriptomic analyses across mouse models and human samples, we describe a gut-metabolic-immune axis that converges on the retina and exacerbates AMD-like degeneration.

Consistent with prior reports that high fat diet alters microbial diversity and promotes expansion of inflammatory taxa (13–15), herein, we provide novel evidence for the impact of a specific diet, with elevated cholesterol burden on gut health. We observed large-scale microbial restructuring accompanied by systemic metabolic signatures of accelerated aging. Amino acid metabolism, particularly histidine pathways, emerged as prominently disrupted. Increase in *Proteobacteria* in the gut has been previously shown to be a key factor for peripheral histidine depletion (42). Also, circulating histidine declines with biological aging in humans (28,29), and multiple metabolomic studies have independently reported reduced histidine and histidine-related metabolites in AMD plasma (68–70). Our findings extend these observations by demonstrating that diet-induced gut microbiome remodeling recapitulates this aging-associated histidine deficiency and links it mechanistically to immune activation. These data support the concept that metabolic insufficiency is not merely correlative but functionally connected to disease progression.

Metabolic stress has been shown to induce trained innate immunity and inflammaging (7,71). In agreement with this framework, histidine depletion in our models was associated with increased levels of inflammatory neutrophils and monocytes, cell populations implicated in AMD pathogenesis (21,65,66,74). Intestinal barrier disruption and activation of IGF1R/AKT2 signaling further align with prior evidence that insulin/IGF-1 pathway regulate longevity, immune tone, and epigenetic remodeling across species (52–54). Our *C. elegans* data demonstrating gut-dependent inflammatory and epigenetic alterations, consistent with reports linking metabolic stress to durable chromatin changes and immune memory in multiple organisms (49–51,53–56). Together, these findings position gut-derived metabolic stress as an upstream driver of systemic immune aging.

In the retina, we demonstrate that systemic histidine deficiency converges with intrinsic RPE vulnerability. SLC7A5 (LAT1), a key transporter of amino acids including histidine, has been implicated in metabolic regulation and mTOR signaling (86). Our observation of reduced SLC7A5 expression in aging and AMD RPE aligns with transcriptomic datasets showing metabolic and inflammatory reprogramming in diseased RPE (87). Our bioinformatic ligand–receptor loop analysis (65) identified MIF/CD74 as a central signaling axis linking metabolically stressed RPE to infiltrating immune cells. MIF is a pleiotropic cytokine elevated in inflammatory and degenerative conditions, and CD74-mediated signaling promotes leukocyte activation and survival (88,89). Prior studies have implicated MIF and CD74 in retinal inflammation (90,91); our data mechanistically connect this pathway to amino acid stress and impaired immune cell clearance by RPE, suggesting a feed-forward inflammatory loop.

Importantly, metabolic stress and immune dysregulation modified disease trajectory and exacerbated RPE damage in a mouse model with an age-dependent chronic dry AMD-like phenotype (65,66,80). Additionally, histidine supplementation, AKT2-PSM therapy, and microbiota restoration attenuated immune activation upon HFC exposure. Further, histidine supplementation and AKT2-PSM therapy rescued retinal degeneration in an acute mouse model of oxidative stress, a known factor in AMD pathobiology (84). Recently, nutrient sensing and amino acid availability are increasingly recognized as modulators of immune and tissue homeostasis (27–30,35–37), and dietary modulation has been proposed as a strategy in AMD prevention (15).

Collectively, our data support a directional model in which diet-induced alterations in gut microbiome functions as the upstream trigger of systemic metabolic deregulation and immune aging. This microbial remodeling reduces circulating histidine, suggesting this as the initiating metabolic event. We identify gut-derived IGF1R/AKT2 signaling as the central regulatory node, integrating metabolic stress with global epigenetic remodeling and innate immune reprogramming. Depletion in peripheral histidine levels leads to expansion and inflammatory priming of peripheral neutrophils and monocytes, generating a state of systemic immune aging. Our study shows that histidine mediated immune regulation converges in the retina, where age-associated downregulation of the major histidine transporter SLC7A5 renders RPE cells metabolically vulnerable. The intersection of systemic immune activation with RPE-intrinsic amino acid transport deficiency indicates a feed-forward inflammatory circuit mediated in part through MIF/CD74 signaling, culminating in chronic inflammation and degeneration. Thus, histidine-axis dysregulation links environmental dietary stress to systemic immune remodeling and tissue-specific pathology through a unified metabolic–immune mechanism.

Notably, while our data identify neutrophils and monocytes and mononuclear phagocytes as the predominant infiltrating populations, we do not determine the retinal involvement of other peripheral immune cells like MSDC, NK or T cell subsets and whether they are also being modulated by the histidine axis. Additionally, to ascertain the histidine metabolic fate in RPE or immune cells, future in-depth studies employing isotope-labeled metabolic flux analysis will need to be performed. Importantly, this study does not exclude the possibility that the HFC diet exerts direct effects on retinal-resident immune cells, such as microglia, or tissue resident macrophages (TRM) independent of gut microbiome, systemic histidine depletion and peripheral immune reprogramming leading to the retinal degenerative changes. Though our results support that gut-mediated alterations in histidine metabolism triggers peripheral immune deregulation and exacerbates retinal inflammation/degeneration, an alternative pathway could be the direct involvement of dietary/metabolic components in regulating retinal immune cells (microglia, TRM). Future experiments and in-depth studies are needed to ascertain the direct impact of diet on retinal immune cells.

## Methods

### Antibodies

The flow cytometry antibodies are CD45-PECy7 (103113, BioLegend), CD11b-Alexa fluor 700 (557960, BD Pharmigen), Ly6C-BV421 (128031, Biolegend), Ly6G-APC (560599, BD Pharmigen), CCR2-FITC (150607, Biolegend), CXCR2-APC/Cy7 (149313, Biolegend), CX3CR1-PerCp-Cy5.5 (149009, Biolegend), F4/80 (12-4801-82, Invitrogen) and CD74 (151002, Biolegend). The secondary antibody used to tag CD74 antibody is Goat anti-rat-Alexa fluor 488 (A-11006, Invitrogen). The antibodies used in immunohistochemistry studies are CCR2 (ab254375, Abcam), E-cadherin (3195S, Cell Signaling), ZO-1 (13663S, Cell Signaling), Rhodopsin (NBP2-25160SS, Novus Biologicals), and EBP50 (PA1-090, Invitrogen). The secondary antibody and cytosolic/nuclear counterstains used for the respective immunohistochemistry studies are Donkey anti-rabbit-Alexa fluor 555 (A31572, Invitrogen), Donkey anti-mouse-Alexa fluor 488 (A21202, Invitrogen), Phalloidin-Alexa fluor 488 (A12379, Invitrogen), Hoechst (62249, Invitrogen) and DAPI (62248, Thermo Fisher).

### Animals

All animal studies were conducted in accordance with the Guide for the Care and Use of Animals (National Academy Press) and were approved by the Johns Hopkins University Animal Care and Use Committee. The authors have complied with all relevant ethical regulations for animal maintenance and euthanasia during this study. RPE-specific βA3/A1-crystallin conditional knockout mice (*Cryba1* cKO) were generated as described previously by our laboratory (80). WT and *Cryba1*-floxed mice were used as controls. All mice that were used in this study were negative for RD8 mutation (80)

### Sex as a biological variable

AMD is affected in both males and females (24), so animals of both sexes were used in our study in equal proportions to ascertain any sex-based bias in experimental results.

### HFC diet

5-month-old C57Bl6/j (WT) mice were fed with either normal chow or HFC diet (TD.88051, Envigo) (67) for 8 weeks; 8-month-old *Cryba1*-floxed and cKO mice were fed with these same diets for 16 weeks and then experiments were performed.

### Human blood isolation

Human venous blood was isolated from donors with or without AMD (Table-1) at the Henderson Ocular Stem Cell Laboratory, Retina Foundation of the Southwest, following their approved institutional protocol (ARMD-SS-2019). Serum isolation and genotyping of each donors for the *CFH* Y402H risk allele was performed using Taqman probe kit (C_8355565_10, Thermo Fisher) (66,78).

### Fecal matter transplantation

WT mice fed with HFC for 4 weeks were subjected to microbiota depletion via oral gavage (200 μL) of equal ratio of 1 g/mL each of an antibiotic cocktail containing neomycin (N5285-25G, Sigma Aldrich), metronidazole (M3761-5G, Sigma Aldrich), ampicillin (A9393-5G, Sigma Aldrich) and gentamycin (G3632-5G, Sigma Aldrich) along with 0.5 g/mL of vancomycin (94747-1G, Sigma Aldrich), administrated daily for 3 consecutive days (92). Immediately after microbiota depletion, fecal sample from normal chow fed WT mice was weighed and suspended in sterile PBS (Thermo Fisher, Cat# 10010-023) and administered to microbiota depleted HFC fed WT mice by oral gavage (300 μL) at a dose of 20 mg/day for 3 consecutive days (93). Animals were euthanized 8 weeks after the initiation of HFC treatment.

### *C. elegans* generation, maintenance and treatments

*Caenorhabditis elegans* strains WT, *Daf2*^ins^ (daf-2(hq363[daf-2::degron::mNeonGreen]) III; MQD2428), and mgEx341 [akt-2::GFP::unc-54 3’UTR + rol-6(su1006)], a transgenic reporter lines were obtained from the CAENORHABDITIS GENETICS CENTER (CGC), University of Minnesota which is funded by NIH Office of Research Infrastructure Programs (P40 OD010440). The worms were maintained on nematode growth medium (NGM) agar plates seeded with *E. coli* OP50 at 25 °C using standard conditions. All experiments were performed on synchronized L4 young-adult worms. To study the effect of high fat (HF) on these worms, OP50 cultures were incubated with 15.8% of lipid mixture (L0288-100ML, Sigma Aldrich) and then spread on NGM plates and incubated for 6 hours at 25°C using standard conditions. Next, to ascertain the involvement of Daf2 signaling in the gut after HF exposure, 1 mM Auxin was used to specifically degrade Daf2 in the intestinal cells of *Daf2*^ins^ worms. Additionally, AKT2-PSM (AKT2i; 10 nM) and histidine (10 mM; H8000-25G, Sigma Aldrich) were also added to selected HF+OP50 cultures and then added to NGM plates before plating the worms.

### Single cell RNA sequencing (scRNAseq) analysis and bioinformatics

RPE-choroid cell suspensions from 3- and 15-month-old WT and *Cryba1* cKO mice were subjected to scRNAseq as described previously (65). Seurat objects for each sample were created using the function “CreateSeuratObject” in Seurat v5.2.1 (94). Cells with nFeature_RNA > 8000, nFeature_RNA < 250, nUMI > 40000, nUMI < 500, log10(GenesPerUMI) < 0.8, or with a mitochondrial rate > 20% were filtered out. Predicted doublets were identified and removed using Scrublet v0.2.1 (95) with the default parameter settings. Then genes detected in less than 5 cells were also removed. As a result, the expression of 17626 genes in 10134 cells were used for downstream analysis. After integration and clustering using Seurat, cell type identities were assigned based on the top marker genes of each cluster as well as visualizing the expression of canonical marker genes of candidate cell types. Average gene expression values in each cell type/cluster and each sample were then calculated by the function “AverageExpression” in Seurat package. Differential expression analysis was performed on each cell type/cluster of interest between cKO and WT samples by the function FindMarkers with test.use = “wilcox” in Seurat package. Gene set enrichment analysis for the KEGG pathways of interest was performed using the function “gseKEGG” of the R package clusterProfiler v4.10.0 (96).

For the analysis of ligand-receptor interactions between cell types/clusters, we used the default built-in ligand-receptor interaction database, intracellular signaling database and gene regulatory networks in the R package LRLoop (65), removed ligand-receptor pairs and intracellular signaling interactions with detection rate less than 2.5% in the corresponding cell types/clusters. The remaining candidate ligand-receptor pairs between each pair of cell types/clusters of interest in each sample were scored by the package LRLoop with its standard pipeline and default parameters. Then for each ligand-receptor pair between each cell type/cluster, the maximum score across WT and cKO samples at each time point was taken as its interaction score for that time point. For the clarity of visualization, the top sixty ligand-receptor pairs of the highest interaction scores among each group of interactions of interest were identified and included in the corresponding Circos plots. For comparative analysis of scRNAseq (mouse) and RNAseq (human iPSC-RPE) differentially expressed genes (DEGs) from mouse and human datasets were identified based on the criteria of the genes having an expression count greater than or equal to the 1.5-fold change (FC) between two experimental conditions and a false discovery rate (FDR) less than 0.05. After respective cutoffs of FC > 1.5, P < 0.05, and FDR < 0.05, the DEGs were identified for each experimental comparison, each list of genes along with their differential expression values was uploaded to Qiagen Ingenuity Pathway Analysis (IPA). Pathway enrichment matrix was represented from z-scores and displayed via heatmap (97). Gene expression analysis was performed from human scRNAseq datasets (87) using the SPECTACLE tool, which is a Single-Cell Atlas of Gene Expression in the Eye, developed by the University of Iowa Institute for Vision Research.

### RNAseq analysis

WT RPE explants (n=3) infected with Ad-RFP, Ad-βA3-RFP, Ad-βA1-RFP and Ad-βA3/A1-RFP constructs (customized from Vector Biolabs) and human iPSC-RPE cells (n=3) from AMD (*CFH* Y402H) and control siblings (*CFH* Y402) donors (78), respectively were subjected to total RNA isolation using a commercially available kit (BIO-52072, Bioline). The isolated RNA was subjected to cDNA library preparation followed by RNAseq and bioinformatic analysis, which was performed as a paid service by Novogene, USA, using their standardized protocols (57).

### 16S rRNAseq and analysis

Fecal matter from *Cryba1*-floxed and cKO mice fed with either normal chow or HFC were harvested from small intestine. Gut bacterial communities of control and HFC diet fed animals were profiled using 16S rRNA sequencing (V4 region). Sample preparation, sequencing and analyses was performed by Novogene following standard protocols. Briefly, paired-end Illumina reads were processed using the QIIME pipelines for quality control of raw data, OTU identification and microbial community analysis. Base R packages, tidyverse and edgeR were used for data visualization.

### Sodium Iodate model and *in vivo* treatment of AKT2-PSM and histidine

The sodium iodate model for dry AMD was generated using previously a published method (84). Briefly, WT mice (5-month-old) were injected intraperitoneally with 20 mg/kg body weight of sodium iodate after 7 weeks of HFC or NC exposure and animals were euthanized after one week (84). Pretreatment of once weekly administration of AKT2-PSM (AKT2i; 0.1 mg/kg, intraperitoneally) or histidine (1.875 g/kg, oral gavage) (98) was given to mice starting from 5th week after HFC exposure and the treatment continued throughout the experimental duration.

### Neutrophil extracellular trap (NET) formation

Human neutrophils (IQB-Hu1-Nu10, IQ Biosciences) were cultured on fibrinogen coated glass bottom plates (66) in presence or absence of 10% human AMD (from *CFH* Y402H donors, Table-1) serum or serum from sibling controls (from *CFH* Y402 donors, Table-1) for 6 hours. To assess the effect of histidine on neutrophil activation, 10 mM of histidine was added to selected AMD serum containing cultures. The neutrophils were fixed and then stained with DAPI (1 μg/ml), the plates were washed with 1× PBS, 5 times, for 15 min each and then mounted with a coverslip using DAKO mounting medium until imaged with Zeiss LSM 880 microscope to assess the formation of NETs in different conditions (66).

### Enzyme-Linked Immunosorbent Assay (ELISA)

ELISA was performed to assess the levels of LCN-2 (ELH-Lipocalin2-1, Ray Biotech) and MPO (ELH-MPO-1, Ray Biotech) in neutrophil lysates from cultures containing 10% human AMD (from *CFH* Y402H donors) serum or serum from sibling controls (from *CFH* Y402 donors) for 6 hours with or without 10 mM histidine. The ELISA was done following the manufacturer’s protocol.

### Flow cytometry

Mouse blood immune cells were analyzed by flow cytometry using previously published methods (61,66). Briefly, blood was collected in heparinized vials and RBCs were lysed using lysis buffer (00-4333-57, Thermo Fisher); whole blood immune cells were harvested as pellet after centrifugation at 1000g for 10 min. Cell pellets were used to assess the changes in immune cell populations by flow cytometry (SONY FACS), after staining with anti-Ly6G, anti-Ly6C, anti-CD11b, anti-CXCR2, anti-CCR2 and anti-CD45 antibodies at a concentration of 1 μg/mL for 90 min at room temperature.

### Metabolomics and histidine quantification

The serum and RPE from different mouse groups (*Cryba1*-floxed and cKO mice fed with either normal chow or HFC) were used for metabolomics analysis (GC-TOF MS) as a paid service from West Coast Metabolomics Center, UC Davis, CA. The metabolomics results were bioinformatically analyzed using Metaboanalyst 6.0. Quantitative assessment of different metabolites was performed at alpha = 0.05 level. For metabolites with significant ANOVA results (p < 0.05), post-hoc pairwise comparisons were conducted using the Bonferroni correction to control the family-wise error rate. The adjusted significance level for pairwise comparisons was $\alpha’ = \alpha/m$, where $m = \binom{4}{2} = 6$ is the number of pairwise comparisons. All analyses were performed using SAS version 9.4 (SAS Institute Inc., Cary, NC). Multiple testing across metabolites was addressed by reporting false discovery rates for significant findings.

Quantitative estimation of histidine levels from human (control and AMD) serum was performed by HPLC as a paid service from the Clinical Pharmacology Analytical Core (CPAC), Indiana University, Indianapolis, IN.

The estimation of histidine from mouse serum was performed quantitatively using sample preparation and analysis methods published previously (99–101) with modifications as described below. Briefly, serum samples were deproteinized using perchloric acid precipitation followed by sequential dilution. Analyte-containing serum was mixed with an equal volume of perchloric acid (1:1, v/v) and vortexed thoroughly to precipitate proteins. The mixture was centrifuged at 14,000–16,000 × g for 5–10 min at 4 °C, and the clear supernatant was carefully transferred to a fresh microcentrifuge tube. The recovered supernatant was subsequently diluted 10-fold with 0.01 M hydrochloric acid to reduce matrix effects and improve chromatographic performance, followed by a second centrifugation under identical conditions to remove any residual particulates. The resulting clarified supernatant was again diluted with 0.01 M hydrochloric acid to achieve a final 10-fold dilution relative to the second-spin supernatant. Before the derivatization and injection, samples were passed through a 0.22 μm syringe or spin filter to protect the chromatographic column. The processed extracts were either analyzed immediately or stored briefly on ice until derivatization.

The pre-column derivatization reagent was prepared fresh daily in sodium borate buffer (pH 9.9). O-phthalaldehyde (OPA, 13.4 mg) was first dissolved completely in 1.0 mL of methanol and then combined with 9.0 mL of sodium borate buffer to obtain a final reagent volume of 10 mL containing 10 mM OPA. Subsequently, 3-mercaptopropionic acid (MPA, 10 μL) was added as the thiol component required for isoindole formation. The solution was mixed gently, filtered through a 0.22 μm membrane, protected from light, and stored at 4 °C. The reagent was used within 24 hours of preparation.

Chromatographic analyses were performed using a Shimadzu Corporation HPLC system equipped with LC-40 series quaternary pumps, a SIL-40C XR autosampler, a CL SIL-40C X3 autosampler module, and both UV–Vis and fluorescence detectors. Instrument control, data acquisition, and peak integration were carried out using Shimadzu LabSolutions software. Separation was achieved on a reversed-phase C18 analytical column using a binary mobile phase consisting of water containing 0.1% (v/v) trifluoroacetic acid (mobile phase A) and acetonitrile containing 0.1% (v/v) trifluoroacetic acid (mobile phase B).

Gradient elution was performed according to the Agilent amino acid analysis program, starting at 2% B (0–0.35 min), increasing linearly to 57% B at 13.4 min, stepped to 100% B at 13.5 min, held until 15.7 min, and returned to 2% B at 15.8 min with re-equilibration until 18.0 min. The flow rate was maintained at 1.5 mL/min, and the injection volume was 25 μL.

OPA-derivatized histidine was detected using fluorescence detection with an excitation wavelength of 347 nm and an emission wavelength of 454 nm. Under these conditions, derivatized histidine eluted at approximately 3.5 min and was quantified based on peak area relative to calibration standards.

### Cryo-sectioning

Mouse retina and gut sections were performed as explained previously (57). The retinal and gut sections slides were incubated with respective primary antibodies prepared in primary antibody buffer (1:100) containing 1% Tween 20 + 0.5% BSA in PBS and incubated in a humidified chamber overnight at 4°C. The slides were washed with PBS + 1% Tween 20 three times. The slides were then incubated at room temperature in the dark for 2 hours with appropriate secondary antibody buffer containing 0.1% Tween 20 + 0.5% BSA in PBS and DAPI/Hoechst or Phalloidin as per the experiments. The slides were washed with 1× PBS, 5 times, for 15 min each and then mounted with a coverslip using DAKO mounting medium until imaged with Zeiss LSM 880 microscope (57).

### RPE flat mount staining

Mouse RPE flat mounts were prepared as explained previously (57). The flat mounts were blocked with appropriate serum (57). Then the flat mounts were stained with Iba1 or ZO-1 (13663S, Cell Signaling Technology) in primary antibody buffer (1% Tween 20 + 0.5% BSA in PBS) and incubated in a humidified chamber overnight at 4°C, after quenching autofluorescence using sodium borohydride (57). After three washes with 1% Tween 20 in PBS, the flat mounts were stained with Anti-rabbit Alexa fluor 555 (1:200; A31572, Invitrogen) and DAPI (1:2000; 62248, Thermo Fisher) in 0.1% Tween 20 + 0.5% BSA in PBS for 2 hours in dark/room temperature (57). The RPE flat mounts were washed 5 times with 1× PBS, 15 min each and then mounted with a coverslip using DAKO mounting medium until imaged with Zeiss LSM 880 microscope and quantification of dysmorphic RPE was performed as explained previously (57).

### RPE mediated immune cell clearance assay

Peripheral immune cells were harvested as explained previously (66). The cells were cultured in presence of MIF (1ng) or amino acid free media (R90100110L, USBiological) with 10% dialyzed fetal bovine serum (A3382001, Thermo Fisher) or both for 6h and then stained with CMTPX red (C34552, Invitrogen) as explained previously (66) and co-cultured with 4-month-old WT RPE explants. The whole RPE explant was stained with Phalloidin-488 and DAPI after three PBS washes the next day and then mounted on a cover slip and visualized under LSM 880 (Zeiss, Switzerland). The number of CMTPX positive cells were counted on RPE surfaces as explained previously (74).

### Quantitative RT-PCR

RNA isolation and cDNA preparation was performed in *C. elegans* using previously published methods (57). Taqman probe (Thermo Fisher) based qPCR was performed using *Mek-2* (Ce02415226_m1) and *Tir-1* (Ce02444988_g1), and *Act3* (Ce02784145_s1) as we have described previously (57).

### DNA methylation

*C. elegans* epigenetic changes were estimated using DNA methylation kit and using the manufacturer’s protocol (P-1030-48, Epigentek).

### Health span estimation in *C. elegans*

Health span assays in nematodes were conducted as illustrated in Figure 4. Animals were placed in a six-well plate illuminated from below by an infrared (IR) LED array (Univivi, USA). The emitted light was diffused through a silicone pad and subsequently through agarose to generate a homogeneous background for bright-field imaging. A varifocal lens (Computar, USA) and a long-pass filter (LPF; Schott, Germany) were mounted on a camera (Teledyne, USA) positioned above the plate.The camera was controlled using custom Python scripts to record 60-second video bursts at 10 Hz every 30 minutes. Each experimental session lasted 24 hours.

Video bursts were processed with a custom Python pipeline implemented in OpenCV (version 4.8.1). A circular region of interest (ROI) was manually defined, and frames were cropped to its bounding box to reduce computational load. Two user-defined lines subdivided the ROI into four quadrants (each treatment group) for regional behavioral analysis. Worm detection was performed independently on each frame. To estimate background intensity, a maximum time-projection was computed across every 100-frames. Each grayscale frame was subtracted from the corresponding background image to generate an absolute-difference image, which was Gaussian-blurred and band-pass thresholded within a user-defined intensity range. Morphological opening with an elliptical structuring element was applied to remove small artifacts, and the resulting binary mask was restricted to the ROI. External contours were extracted and filtered by area to exclude objects outside the expected size range of a worm. For each retained contour, the enclosed binary mask was skeletonized using Zhang–Suen thinning to obtain the medial axis. The longest geodesic path within the skeleton was identified, and seven equally spaced points were interpolated along this path to generate an ordered spine representation.

Detections were linked across consecutive frames to maintain persistent animal identities. Centroid coordinates were used to compute a pairwise Euclidean distance matrix between frames, and optimal assignments were determined using the Hungarian algorithm with a maximum allowable displacement of 50 pixels. Detections exceeding this threshold were not matched. Unmatched contours were assigned new identities, and tracks were terminated if no match was found in subsequent frames. Head-tail orientation was inferred from whole-contour motion. For each tracked animal, filled contour masks from consecutive frames were compared to identify advance regions (pixels newly occupied) and retreat regions (pixels vacated). The vector from the centroid of the retreat region to that of the advance region represented net translational motion of the body. This vector was projected onto the spine axis to determine directional consistency with the current head assignment. Evidence for or against the assigned orientation accumulated across frames. The spine orientation was reversed only after three consecutive frames of contrary evidence, after which the evidence score was reset with a positive bias to introduce hysteresis and prevent rapid oscillation during pauses or low-displacement periods. The head-bending angle was computed on the oriented spine as the angle between the first two segments at the head end (102).

Two animal-level exclusion criteria were applied prior to statistical summarization. First, animal identities persisting for fewer than five frames were discarded as transient segmentation noise. Second, identities with zero cumulative centroid displacement were excluded as stationary debris. For each remaining animal, mean head-bending angle across valid frames. Per-animal metrics were aggregated by region and burst (time point). For each region at each time point, the arithmetic means and standard error across all included animals were calculated for head-bending angle.

### Statistics

Analyses were performed using the GraphPad 8.0 software and Microsoft Excel. Differences between multiple groups were analyzed using one-way ANOVA followed by Tukey’s post-hoc test and differences between only two groups were estimated by Student’s t-test (57,65,66). Differences were considered significant when p-value<0.05. The analyses were performed on at least three biological replicates (n). All the values are presented as mean ± standard error of the mean (SEM) (57,65,66).

### Study approval

The mouse studies were approved by the Institutional Animal Care and Use Committee (IACUC) of The Johns Hopkins University School of Medicine (protocol #MO24M177), while the human studies were conducted at the Retina Foundation of the Southwest under their approved protocol #ARMD-SS-2019.

## Data availability

All sequencing data will be uploaded to the GEO NCBI database upon acceptance.

## Author Contributions

S.G. and D.S. conceived, designed and supervised the study and secured funding. S.G., V.K., S.B., V.S.B., J.N., and P.D. performed experimental studies. Y.X., A.K.M, and J.Q. provided bioinformatics expertise. S.G., D.M., H.W., J.Y. performed *C. elegans* health span study microscopy and analysis. A.S., A.I.A., and P.P. performed histidine quantification from human plasma. J.D. and D.B. performed statistical analysis on the metabolomics dataset. D.C. S.A.M. and R.M.K. performed and analyzed mouse serum estimation of histidine. S.R.S. generated and characterized iPSC-RPE lines and provided human plasma samples for the study. S.G. and D.S. interpreted the results. S.G. and D.S. co-wrote the manuscript. S.H., S.R., S.G., J.T.H. and D.S. reviewed and edited the manuscript. All authors have approved the final version.

## Acknowledgements

This study was supported by NIH K99EY033421 (S.G.), Foundation Fighting Blindness- Free Family AMD Research Award BR-CMM-0424-0882-TUF (D.S. & S.R.), The Johns Hopkins University School of Medicine start-up funds (DS), funds from the Frieda Derdeyn Bambas Professorship in Ophthalmology (D.S.), P30EY001765 core award from the National Eye Institute, NIH to the Wilmer Eye Institute, The Johns Hopkins University School of Medicine, and unrestricted funds from Research to Prevent Blindness Inc., NY to the Wilmer Eye Institute, The Johns Hopkins University School of Medicine. The authors would like to acknowledge Dr. J. Samuel Zigler., Jr. for reading and providing necessary edits to the manuscript.

**<u>Movie-1</u>**: ***C. elegans* health span.** Movie showing movement of *C. elegans* in chamber explained in Figure 4 with untreated, HF-exposed, HF+ AKT2-PSM (AKT2i) and HF+histidine exposed *Daf2*^ins^ worms.

## Notes

Conflict of interest statement: S.G., V.S.B., S.B., S.H. and D.S. declare a competing interest for U.S. patent application #63/902,825 which has been filed.

