## Supplementary Figures and Legends for "Gut-derived metabolic reprogramming drives immune aging and tissue degeneration"

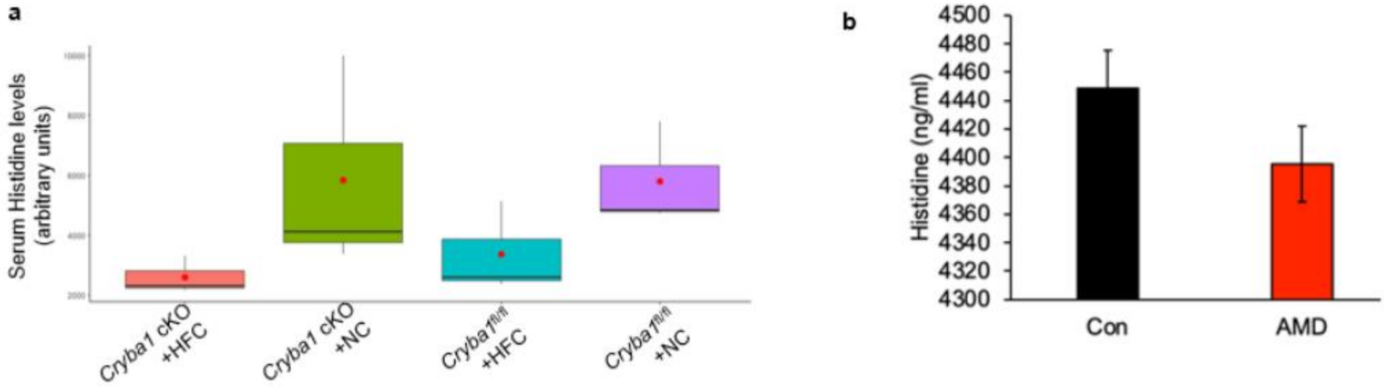


**<u>Supplementary Figure 1</u>**: **Histidine levels in mouse and human blood**. (**a**) Box plot showing significantly (ANOVA, P<0.05) reduced levels of histidine from metabolomics analysis on serum from HFC-fed *Cryba1*-floxed and cKO mice, compared to serum from normal chow fed mice. n=4. (**b**) Quantitative assessment of histidine from human serum samples showed a relative decrease in the histidine level in AMD serum, compared to controls (Control; n=4, AMD; n=7).

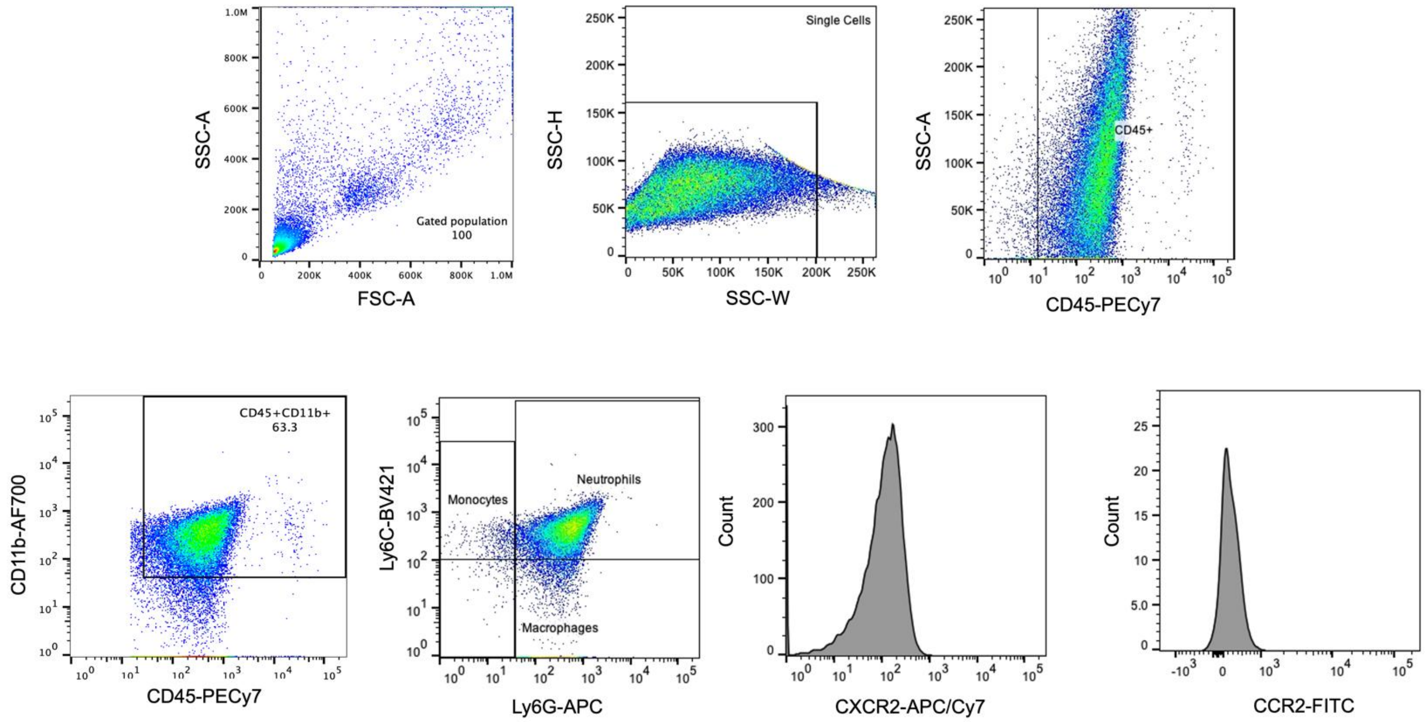


**<u>Supplementary Figure 2</u>**: **Gating strategy for peripheral immune cells**. Peripheral immune cells from NC or HFC fed mice were analyzed by flow cytometry; the % neutrophils, % monocytes and the expression of CXCR2 in neutrophils and CCR2 in monocytes were evaluated. n=4.

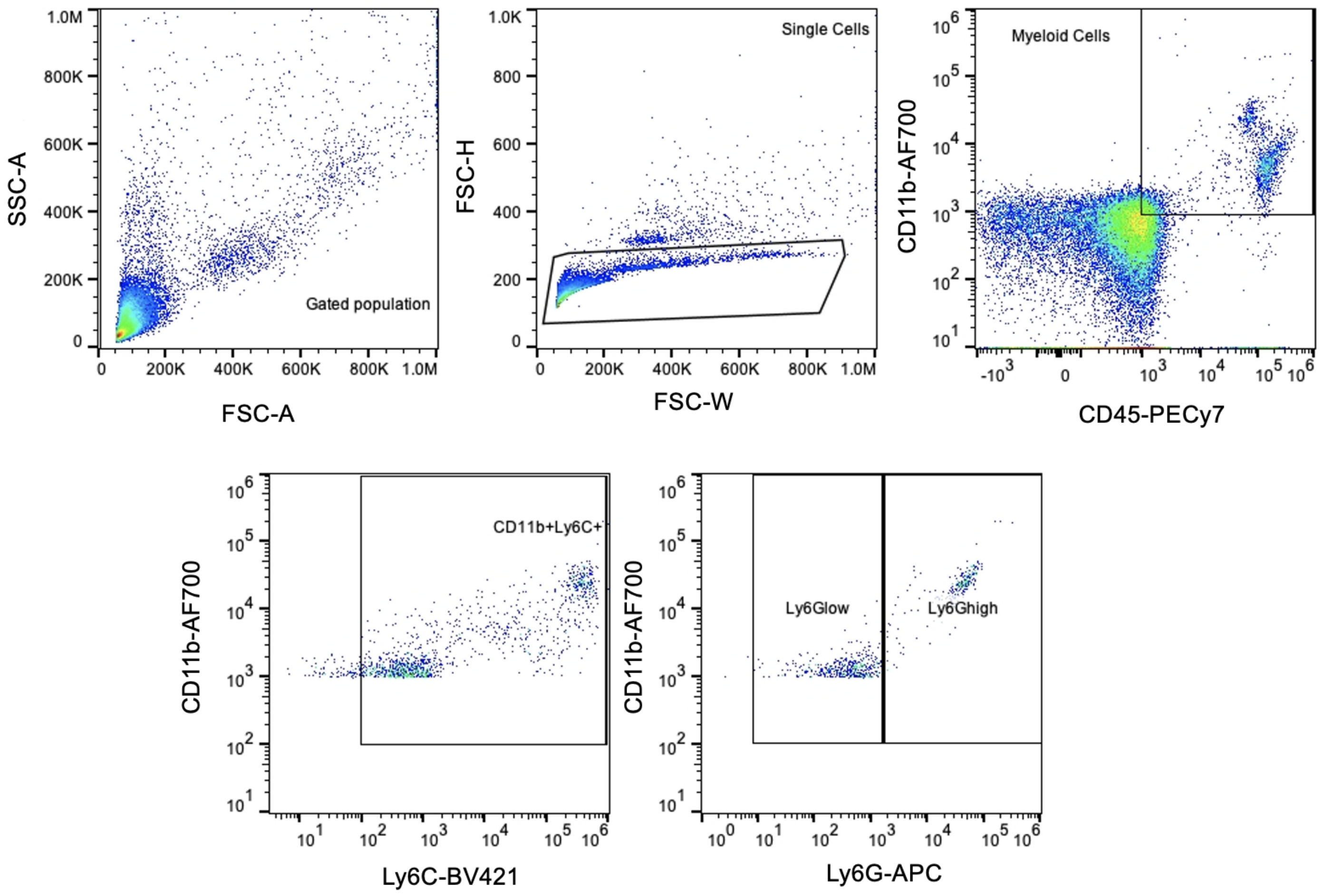


**<u>Supplementary Figure 3</u>: Gating strategy for peripheral immune cells with or without fecal matter transfer**. Peripheral immune cells from NC or HFC fed mice were analyzed by flow cytometry and the %$CD45^{high}CD11b^{high}Ly6C^{+}Ly6G^{high}$ (neutrophils), %$CD45^{high}CD11b^{high}Ly6C^{+}Ly6G^{low}$ (inflammatory monocytes or M-MDSCs) were evaluated. n=3.

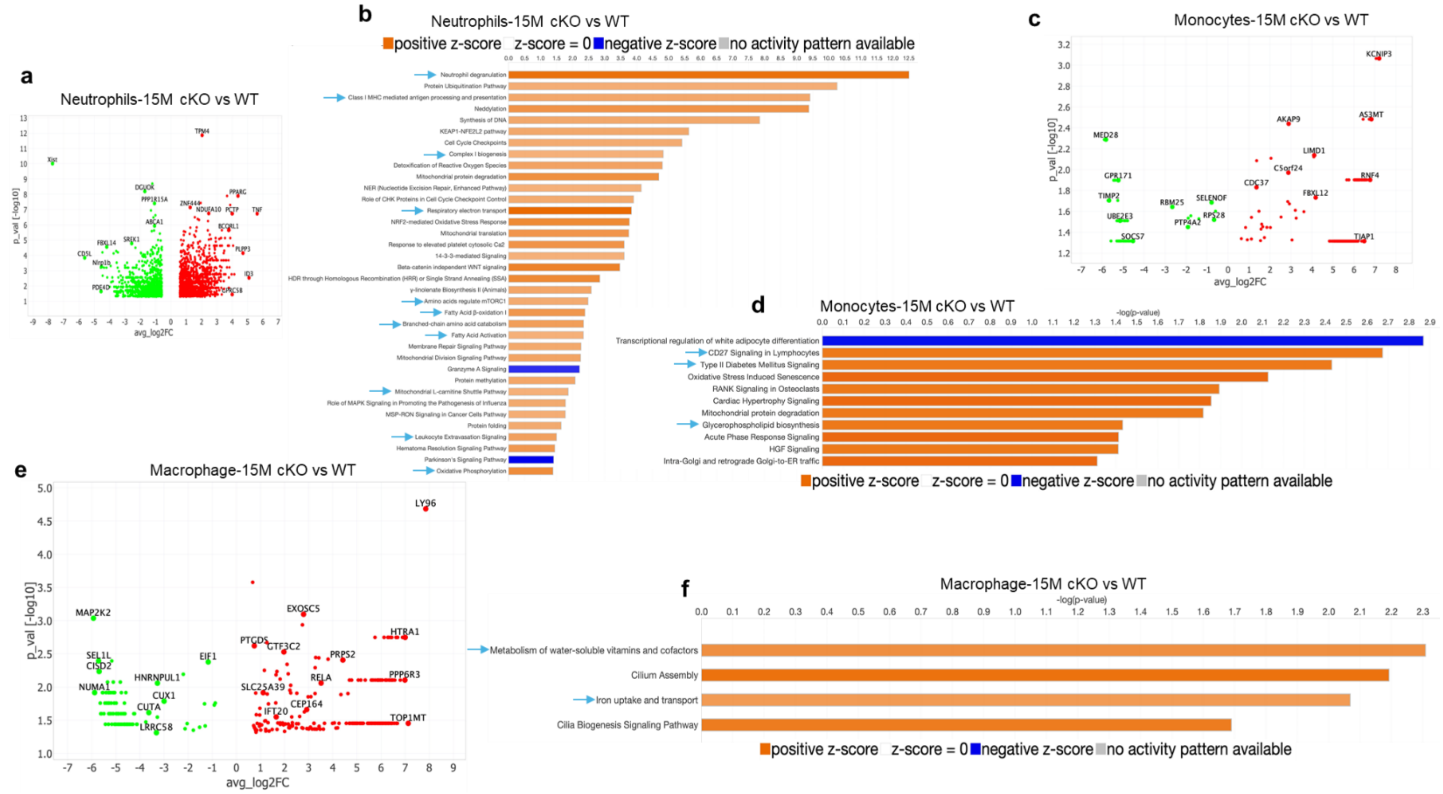


**<u>Supplementary Figure 4</u>**: **Gene and pathway enrichment in infiltrating immune cells**. scRNAseq analysis from 15-month-old *Cryba1* cKO vs WT RPE/choroid tissue revealed noticeable enrichment in genes and pathways that regulate immune homeostasis, metabolism and pro-inflammatory mediators (arrows) in (**a**,**b**) neutrophils, (**c**,**d**) monocytes and (**e**,**f**) macrophages. n=3.

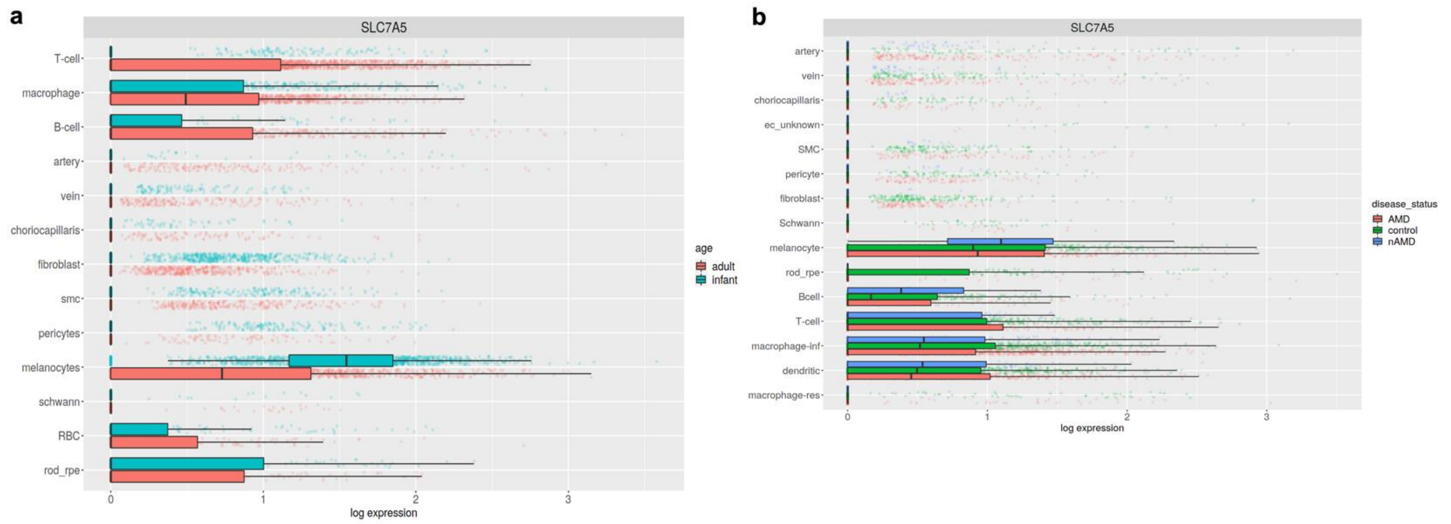


<u>**Supplementary Figure 5**</u>: **Expression of *Slc7a5* in human RPE**. Spectacle software generated representation of different retinal cell population from previously published (87) human scRNAseq analysis revealed a downregulation trend in *Slc7a5* gene expression in the RPE cells (rod_rpe cluster) (**a**) with increasing age [Adult (ages 66-93); n=6, infant (liveborn, ages 1day-9 months); n=2] and (**b**) in neovascular (nAMD) and non-neovascular/atrophic AMD donors, compared to age-matched controls. Additionally, inflammatory macrophages (macrophage-inf) also showed similar trend in AMD tissues (**b**) (Control; n=11, neovascular AMD; n=2, non-neovascular/atrophic AMD; n=9).

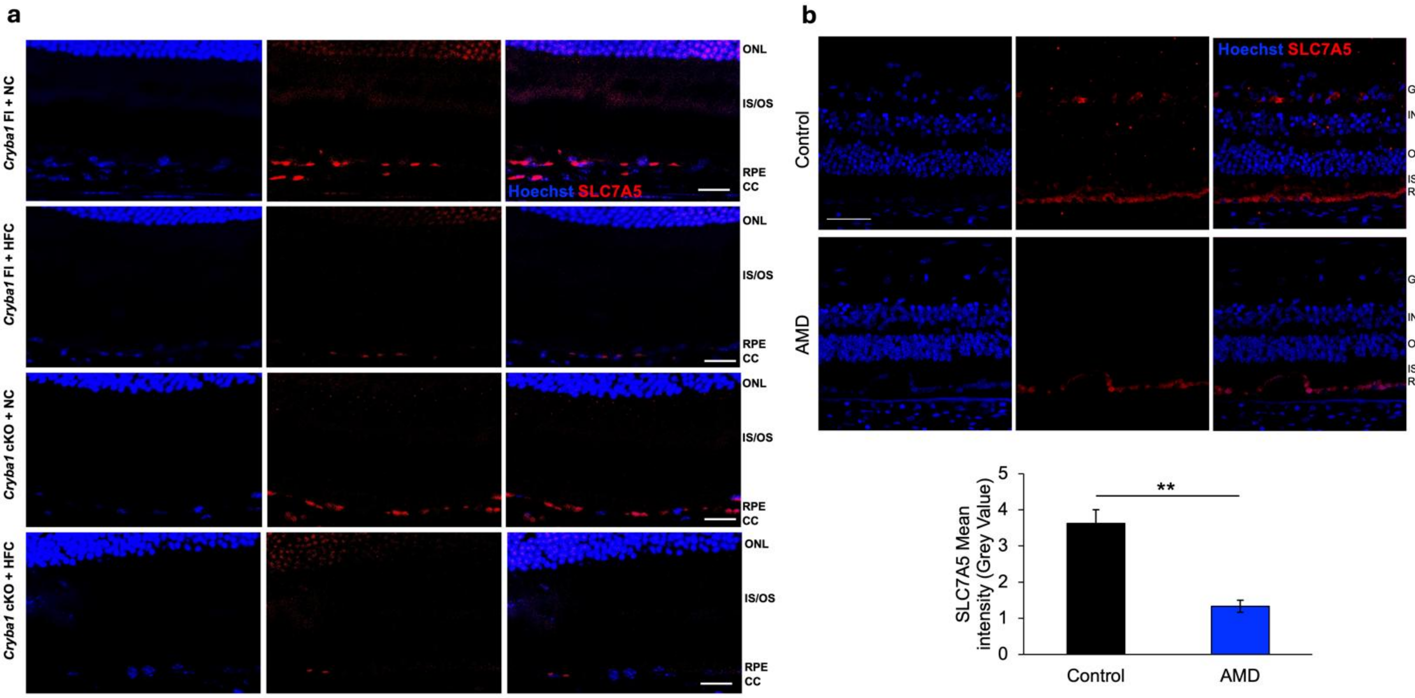


**<u>Supplementary Figure 6</u>**: **Retinal expression of SLC7A5**. Immunohistochemistry analysis showed noticeable decrease in SLC7A5 (red) expression in the RPE of (**a**) *Cryba1*-floxed and cKO mice upon HFC exposure. Normal chow fed cKO mice also showed relative decrease in the expression compared to floxed. n=3. (**b**) Significant decrease in the protein expression was also observed in the RPE of human dry AMD sections. n=3. *P<0.01.

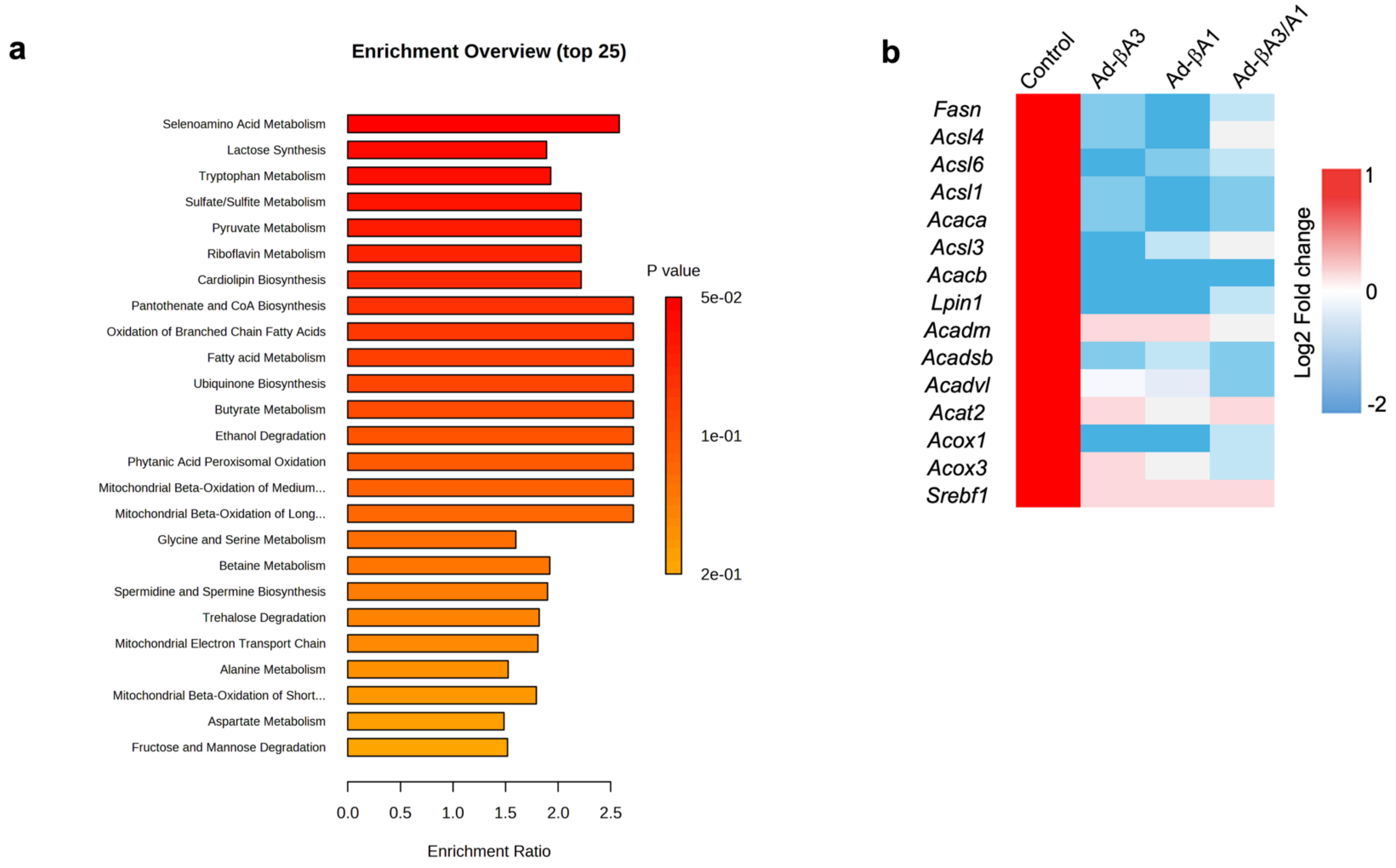


<u>**Supplementary Figure 7**</u>: ***Cryba1* is a key regulator of RPE metabolism.** (**a**) Over-representation analysis bar plot from RPE metabolomics (GC-TOF MS) analysis from 4-month-old *Cryba1*-floxed and cKO mice, showing enrichment of several amino acid metabolism pathways. n=5. (**b**) Heatmap showing expression of major lipid metabolism genes in WT RPE explants overexpressing the native protein or the two crystallin isoforms. n=3.

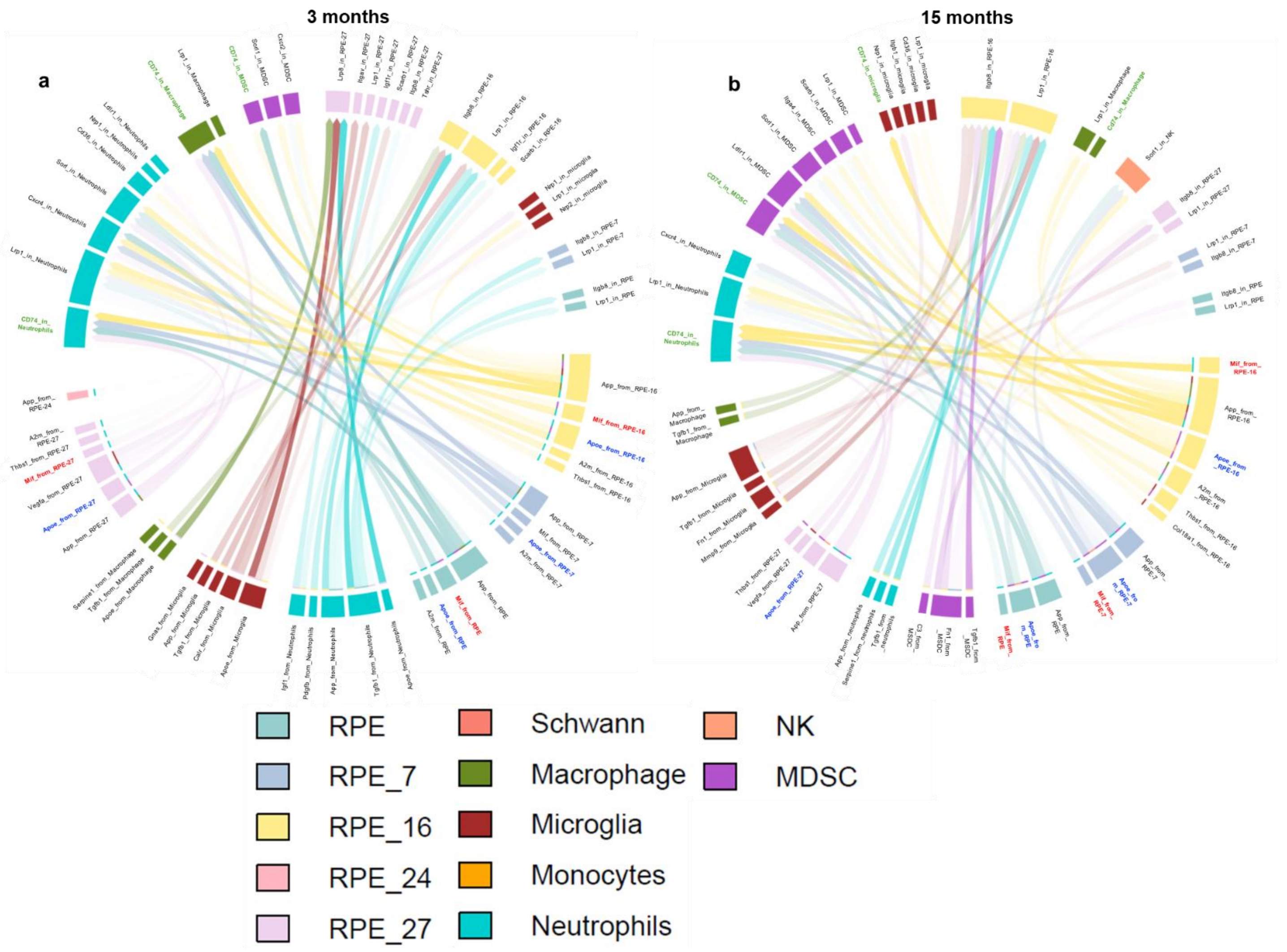


**<u>Supplementary Figure 8</u>**: **LR-loop interaction**. scRNAseq analysis derived interactions between different RPE clusters and immune cells at (**a**) 3- and (**b**) 15 months of age in *Cryba1* cKO vs WT RPE/choroid tissue. The *Slc7a5* downregulated RPE subpopulations/clusters (RPE_16 and RPE_27) showed noticeable expression of several inflammatory molecules like MIF (Macrophage Migration Inhibitory Factor; red), APP (Amyloid Precursor Protein), APOE (Apolipoprotein E; blue), A2M (Alpha-2-Macroglobulin), and THBS1 (Thrombospondin-1), while infiltrating immune cells (neutrophils and macrophages) expressed their cognate receptors, with strong interactions observed between MIF/CD74 (green) and APOE/CD74. n=3.

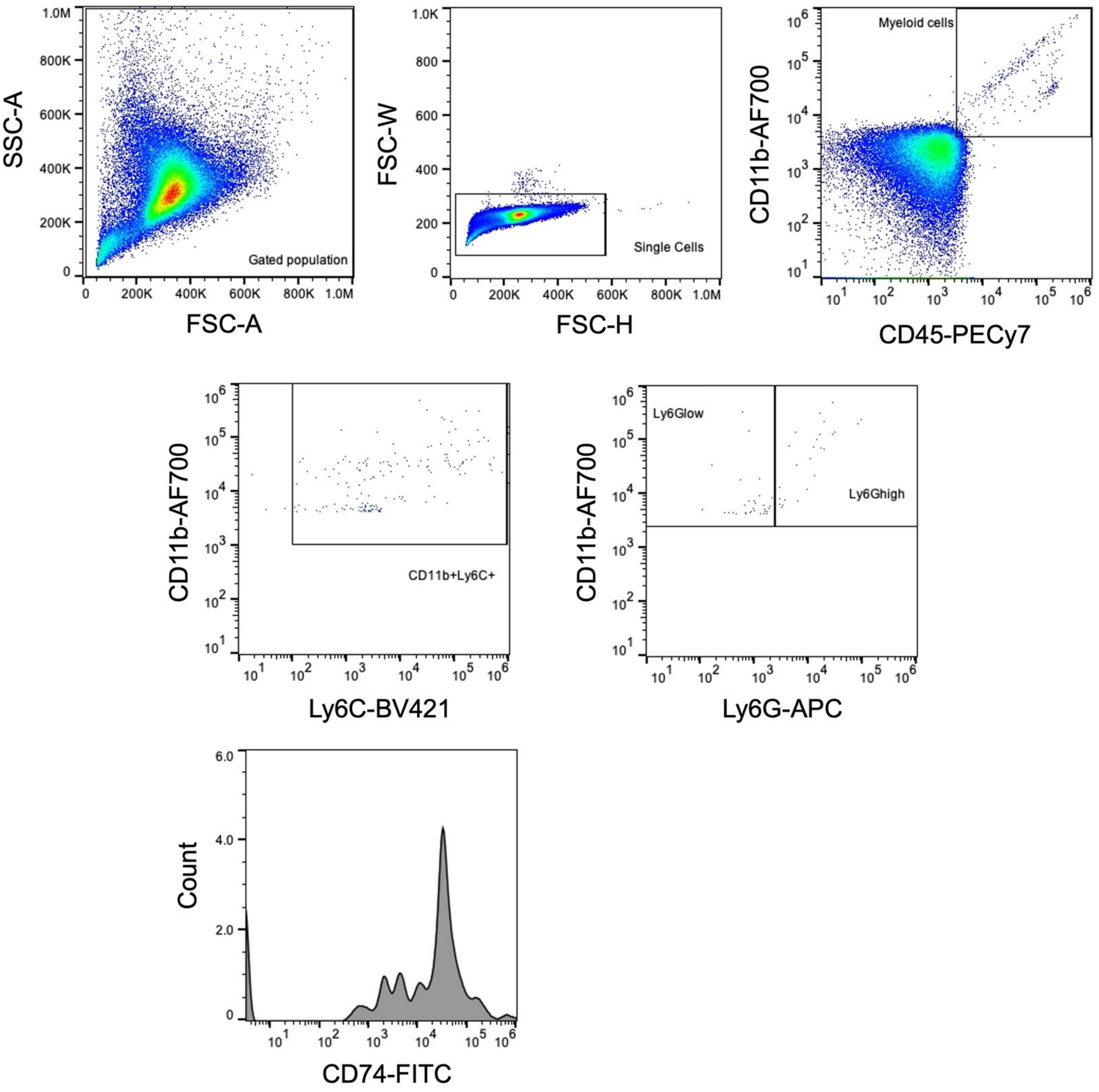


**<u>Supplementary Figure 9</u>: Gating strategy for CD74 expression in cultured peripheral immune cells**. Peripheral immune cells cultured with either MIF protein or amino acid free media or MIF+amino acid free media for 6 hours were analysed by flow cytometry and the expression of CD74 was evaluated in %CD45$^{high}$CD11b$^{high}$Ly6C$^{+}$Ly6G$^{high}$ (neutrophils). n=3.

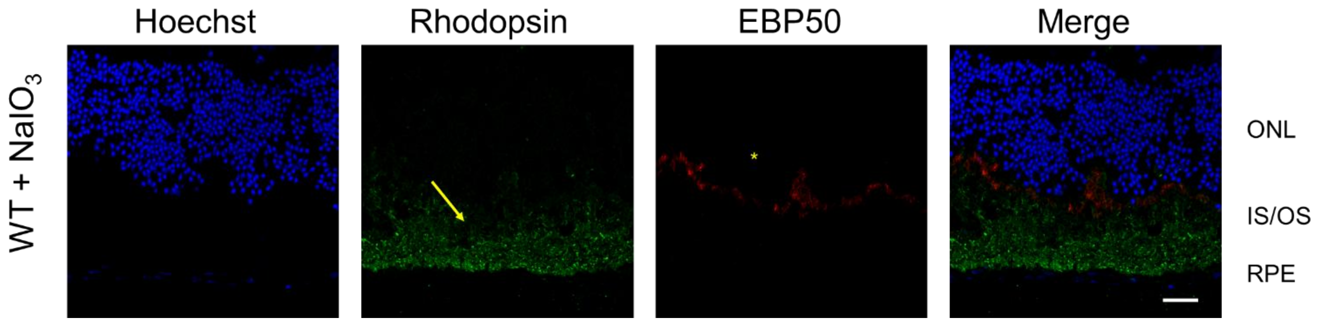


**<u>Supplementary Figure 10</u>**: **NaIO3 model**. Retinal immunofluorescence analyses showing decline in rhodopsin (green; arrow) and EBP50 levels (red; asterisk) compared to WT (as described in Fig.6c). Scale bar= 20 μm. n=3.

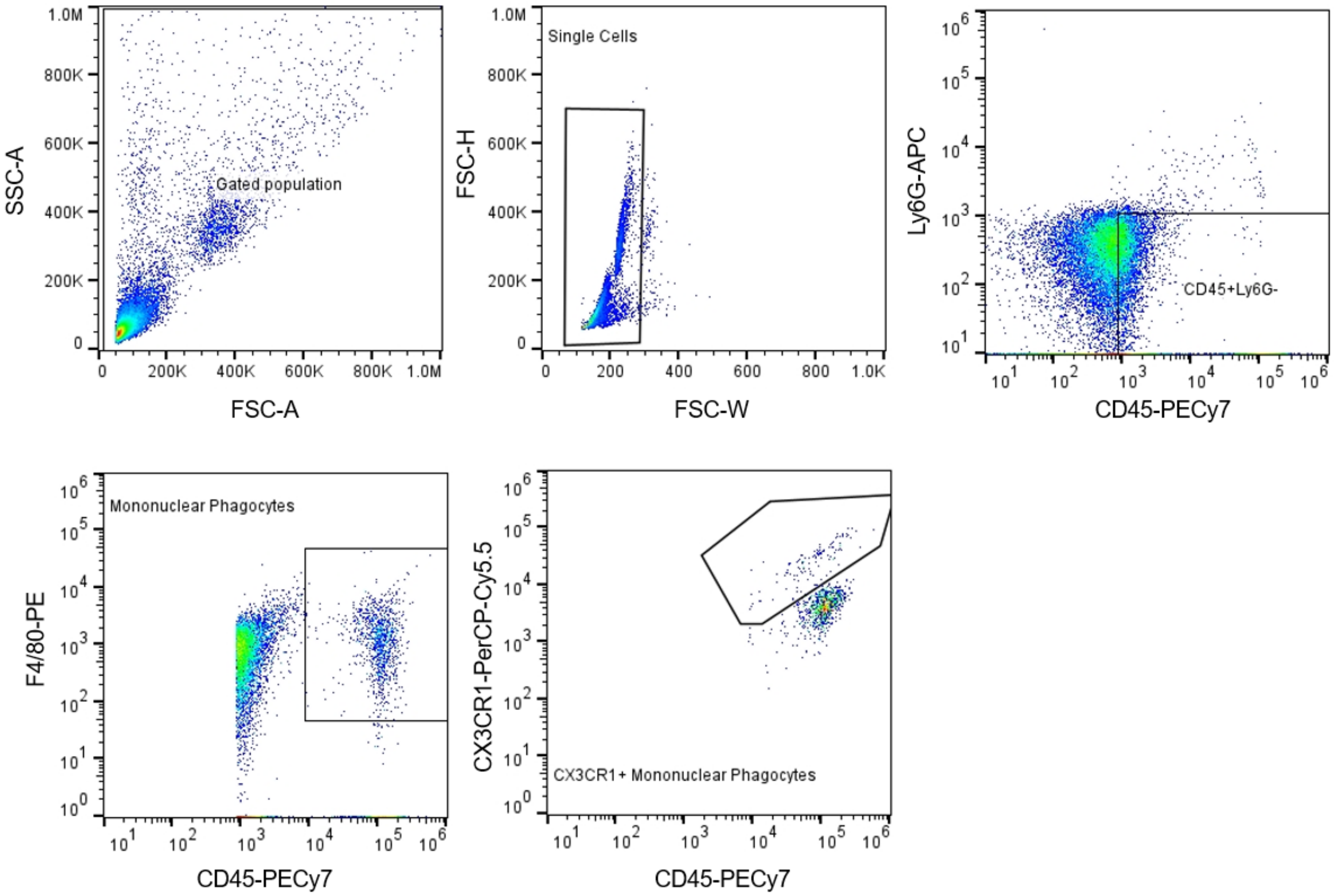


<u>**Supplementary Figure 11**</u>: **Gating strategy for CX3CR1$^{+}$ mononuclear phagocytes among peripheral immune cells**. Peripheral immune cells were analyzed by flow cytometry and the levels of CX3CR1$^{+}$ mononuclear phagocytes (CD45$^{high}$Ly6G$^{-}$F4/80$^{+}$) was evaluated. n=3.

| | | Height | | | Weight | | |
|---|---|---|---|---|---|---|---|
| Patient ID | Disease | Feet | Inches | Cm | Lbs | Kgs | BMI ($kg/m^2$) |
| RFSC#3 | Control | 5 | 5 | 165.1 | 155 | 70.30676 | 25.8 |
| RFSC#4 | AMD | 5 | 5.75 | 167.005 | 143 | 64.86366 | 23.3 |
| RFSC#7 | Control | 6 | 6 | 198.12 | 150 | 68.0388 | 17.3 |
| RFSC#9 | AMD | 5 | 8 | 172.72 | 176 | 79.83219 | 26.8 |
| RFSC#18 | Control | 6 | 0 | 182.88 | 205 | 92.98636 | 27.8 |
| RFSC#19 | AMD | 5 | 10 | 177.8 | 151 | 68.49239 | 21.7 |
| RFSC#22 | Control | 5 | 7 | 170.18 | 140 | 63.50288 | 21.9 |
| RFSC#27 | AMD | 5 | 4 | 162.56 | 174 | 78.92501 | 29.9 |
| RFSC#28 | AMD | 5 | 4 | 162.56 | 160 | 72.57472 | 27.5 |
| RFSC#25 | AMD | 5 | 5 | 165.1 | 115 | 52.16308 | 19.1 |
| RFSC#23 | AMD | 5 | 8 | 172.72 | 150 | 68.0388 | 22.8 |

**Table-1**: **Control and AMD donor details.** Serum isolation was performed from the donors for histidine estimation and human neutrophil culture experiments. Human donors were categorized according to Minnesota Grading System; the AMD donors were MGS2 (dry AMD), whereas controls were MGS1. The table also shows BMI calculations for each donor.
